# Unraveling the Genomic Diversity and Admixture History of Captive Tigers in the United States

**DOI:** 10.1101/2023.06.19.545608

**Authors:** Ellie E. Armstrong, Jazlyn A. Mooney, Katherine A. Solari, Bernard Y. Kim, Gregory S. Barsh, Victoria B. Grant, Gili Greenbaum, Christopher B. Kaelin, Katya Panchenko, Joseph K. Pickrell, Noah Rosenberg, Oliver A. Ryder, Tsuya Yokoyama, Uma Ramakrishnan, Dmitri A. Petrov, Elizabeth A. Hadly

**Author notes:** Contributed equally (Ellie E. Armstrong, Jazlyn A. Mooney).

## Abstract

Genomic studies of rare and endangered species have focused broadly on describing diversity patterns and resolving phylogenetic relationships, with the overarching goal of informing conservation efforts. However, few studies have investigated the genomic diversity potentially housed in captive populations. For tigers (*Panthera tigris*) in particular, captive individuals vastly outnumber those in the wild, yet the diversity of the captive population remains largely unexplored. Here, we present the first large-scale genetic study of the private (non-zoo) captive tiger population in the United States (U.S.), also known as ‘Generic’ tigers. We find that the U.S. Generic tiger population has an admixture fingerprint comprising all six extant wild tiger subspecies (*P. t. altaica*, Amur; *P. t. tigris*, Bengal; *P. t. corbetti*, Indochinese; *P. t. jacksoni*, Malayan; *P. t. amoyensis*, South China; *P. t. sumatrae*, Sumatran). We show that the Generic tiger population has a comparable amount of genetic diversity to most wild subspecies, relatively few private variants, and fewer deleterious mutations. We also observe inbreeding coefficients that are similar to wild populations, suggesting that inbreeding in captive populations is not pervasive, although there are some individuals within the Generic population that are substantially inbred. Our results elucidate the admixture history of the Generic tiger population in the U.S. Additionally, we develop a reference panel for tigers and show that it can be used with imputation to accurately distinguish individuals and assign ancestry even with ultra-low coverage (0.25×) data. The study and reference panel will provide a resource to assist in tiger conservation efforts.

## Main

The Anthropocene has been characterized by population declines, isolation, and extinction, leading to a shift in the abundance and distribution of thousands of species on Earth (1). The tiger (*Panthera tigris*) is one of the most iconic and captivating terrestrial species on the planet and exemplifies the severe and rapid declines in population size and reduced connectivity that threatens the survival of many species (2). Although some tiger populations and subspecies have boasted recent recovery efforts, others have gone extinct in the wild (3) or have been completely extirpated (4) due to human pressures. Today, tigers are composed of six subspecies (*P. t. altaica*, Amur; *P. t. tigris*, Bengal; *P. t. corbetti*, Indochinese; *P. t. jacksoni*, Malayan; *P. t. amoyensis*, South China; *P. t. sumatrae*, Sumatran) that have been genetically and geographically separated for at least the last 10,000 years (5, 6). Their geographic ranges span the Russian Far East and northeast China (*P. t. altaica*), Bangladesh, Bhutan, India, and Nepal (*P. t. tigris*), Myanmar, Thailand, and Laos (*P. t. corbetti*), the island of Sumatra (*P. t. sumatrae*), and the Malay Peninsula (*P. t. jacksoni*), respectively, while the South China tiger (*P. t. amoyensis*) is extinct in the wild with the last sighting more than 30 years ago (7). All subspecies have undergone severe population bottlenecks over the last century, primarily due to human activities such as hunting and the expansion of agriculture, which have directly reduced tiger numbers, habitat availability, and prey (8).

In contrast to wild populations, the global captive tiger population is now estimated to include 15,000–20,000 individuals worldwide, a number that is at least five times larger than the wild population (7). Most populations are largely unregulated, such as farmed and other privately owned tigers, while others, such as those in accredited zoos, are bred with the intention of serving as a diversity reservoir for dwindling wild populations. In the United States, the Association of Zoos and Aquariums (AZA), manages several tiger populations as distinct subspecies, specifically the Amur (1950s-present), Sumatran (1950s-present), Malayan (1980s-present), and for a time the “Bengal” (white tigers; 1960s-2011) tiger subspecies.

The idea that captive populations may serve as diversity reservoirs is a persistent theme and justification of the existence of captive populations (9). For example, the regulated Amur tiger captive population was established in the early 1950s and has since increased to over 1,400 individuals in captivity worldwide (AZA; 10). Previous work which investigated diversity levels of 12 microsatellites in the Amur captive population in North America found that the population may be a reservoir of genetic diversity which is now extinct in the wild (11). This notion suggests that captive breeding programs with effective management are able to maintain genetic diversity and avoid inbreeding depression despite the small number of founders (in the case of the Amur tigers, 29 males and 28 females; (10). Genomic perspectives on whether this is true for many captive populations are rare and the conclusions are inconsistent across species (12–17).

Though there are a large number of tigers in captivity, only a small fraction of these tigers (less than 1%) reside in regulated or accredited facilities, such as accredited zoos (7). Outside of regulated establishments, individuals and entities breed, own, and sell tigers both legally and illegally. The formation of the privately-owned, captive population in the U.S. likely coincided with the establishment of zoos and circuses in the early 1900s, since these entities were known to exchange individuals (8), but ultimately the ancestry and establishment timing of the captive, privately-owned population is unknown. Over time, the mission of AZA zoos began to run counter to circuses and other private ownership practices, but by the 2000s, estimates put the captive tiger population in the United States at greater than 5,000 individuals (8, 18). Many of the unregulated facilities do not report basic or reliable population data and have emerged as a concern globally with the illicit farming of big cats for profit being exposed in Thailand (19), South Africa (20), and most recently the United States, which was popularized by the release of Netflix’s *Tiger King*. In the United States, up until the recent passing of federal legislation (21; Big Cat Public Safety Act), there was very little oversight over the breeding, owning, and selling of tigers.

The pace of breeding in privately owned tiger populations has prompted concerns of inbreeding, especially since phenotypes (i.e. striped whites, snow whites, and golden tigers) that rely on rare, recessive mutations (22, 23) are considered to be of high economic value. In fact, the white tiger (“Bengal”) breeding program was discontinued by the AZA when the intentional selection for the single variant responsible for producing the white phenotype (22, 23) (occurring naturally, but rarely, in wild Bengal tiger populations) was determined to not be relevant to conservation initiatives. This population was also intentionally mixed with Amur tigers in an attempt to prevent inbreeding depression before the program was ultimately discontinued (8). A previous study documented additional signatures of admixture in captive, privately-owned tigers from various localities using mitochondrial DNA (mtDNA) and microsatellites, and suggested that captive tigers contain unique diversity compared to their wild and captive single-ancestry counterparts (24). A separate study also posited that the captive Amur population may contain diversity now extinct in the wild (11). However, neither of the previous studies examined the extent of inbreeding or genetic load in the population using genomic data, which has significantly more power than typical microsatellite datasets to answer such questions. It remains unclear how or if this large population of privately-owned, captive tigers could possibly fit into current management or conservation plans, and whether their genomes hold relic or unique diversity, or on the other hand, if they show indications of severe inbreeding or high genetic load.

For the purposes of conservation genomics, the most useful definition of genetic load is one that describes a reduction in fitness at the population level due to deleterious mutations (25). The genetic load of a population can affect overall population health and viability, and can be modified by stochastic processes such as drift, admixture, and inbreeding (26, 27). Recent genomic-scale work in wild populations has shown purging of deleterious mutations in some groups (28–30), while other groups (31, 32) appear to have a large fraction of deleterious mutations remaining. In fact, recent work in wild Bengal tigers has shown purging of deleterious variation in smaller populations relative to larger populations, alongside continued inbreeding depression due to high frequency deleterious alleles (33). However, there is a dearth of literature on genetic load and inbreeding in captive tiger populations, or other tiger subspecies, at the genomic scale. As genetic rescue, translocation, and captive breeding programs are being implemented more frequently in endangered species for strategic management (34, 35), a broader understanding of genetic diversity and genetic load within captive and wild tiger populations warrants further investigation.

Here, we use genomic data to examine admixture and population structure, quantify genetic diversity and genetic load, and investigate the extent of inbreeding in the captive, privately-owned tiger population in the United States (also known as ‘Generic’ tigers) using whole-genome sequence data obtained from individuals in accredited sanctuaries. These sanctuaries do not breed, sell, or buy tigers, but house tigers previously rescued from unregulated facilities or those that have been forfeited by private owners. We combine our newly sequenced individuals with previously published data, resulting in a dataset representing 255 unique individuals across all six extant wild subspecies and U.S. Generic tigers (Table 1). These data allow us to determine how admixture events have shaped the genomic landscape of the privately-owned captive population. We compare the Generic tigers to their potential source populations (tigers of single ancestry, i.e., subspecies), and examine how diversity is partitioned across each group. Then, we test for potential signs of inbreeding and quantify the total amount of deleterious variation in each population. Lastly, we show that ultra-low coverage (0.25×) data can be imputed sufficiently using reference haplotypes from single ancestry tigers to determine ancestry and perform individual identification. Our results demonstrate that low-coverage sequencing and imputation provide a simple and cost-effective alternative compared to microsatellites and custom SNP panels to identify the source populations and identity of illegally traded individuals or wildlife materials from tigers and for long-term monitoring of captive and wild populations.

**Table 1.** Number of individuals (duplicates removed) used in each analysis by method (imputed/unimputed) with brief description of filters used. See Methods and Supplementary Methods for a additional and specific details.

| Group | Total Individuals |  | Total | PCA | Admixture | Heterozygosity (unimputed, <20% missing) | ROH, IBD, Genetic Load (unrelated, >5K) | SFS (unrelated, >5K, even sampling) | Local Ancestry (unimputed) |
| --- | --- | --- | --- | --- | --- | --- | --- | --- | --- |
| Amur | 38 | Unimputed | 32 | 32 | 32 | 28 | 22 | 10 & 5 | 32 |
|  |  | Imputed | 6 | 6 | 6 |  |  |  |  |
| Bengal | 23 | Unimputed | 23 | 23 | 23 | 23 | 21 | 10 & 5 | 23 |
|  |  | Imputed | 0 | 0 | 0 |  |  |  |  |
| Indochinese | 6 | Unimputed | 6 | 6 | 6 | 6 | 6 | 6 | 6 |
|  |  | Imputed | 0 | 0 | 0 |  |  |  |  |
| Malayan | 22 | Unimputed | 22 | 22 | 22 | 22 | 20 | 10 & 5 | 22 |
|  |  | Imputed | 0 | 0 | 0 |  |  |  |  |
| South China | 4 | Unimputed | 1 | 1 | 1 | 1 | 1 |  |  |
|  |  | Imputed | 3 | 3 | 3 |  |  |  |  |
| Sumatran | 18 | Unimputed | 18 | 18 | 18 | 18 | 15 | 10 & 5 | 18 |
|  |  | Imputed | 0 | 0 | 0 |  |  |  |  |
| Generic | 144 | Unimputed | 68 | 68 | 68 | 50 | 16 | 10 & 5 | 68 |
|  |  | Imputed | 76 | 76 | 76 |  |  |  |  |

## Results

### Population structure and ancestry in Generic tigers

Using both imputed and unimputed individuals, we investigated population structure and ancestry of the Generic tigers using a combination of principal component analysis (PCA) and supervised ADMIXTURE. Both the supervised ADMIXTURE analysis (Fig. 1A and *Supplementary Notes*) and PCA (Supplementary Fig. S1), show that Generic tigers are admixed between the six extant tiger subspecies, with individuals varying in the proportion of their genome derived from each ancestral population (Fig. 1A). Investigation of mitochondrial haplotypes revealed clustering with all but the Sumatran and South China haplotypes (Supplementary Fig. S2-4). Local ancestry analyses (unimputed samples only) revealed tracts from all subspecies except South China, which was not included due to its sample size (N = 1; Supplementary Fig. S5). Overall, we found that none of the Generic tigers tested have single-subspecies ancestry. Generic individuals, on average, are primarily admixed between the Amur and Bengal tiger subspecies (mean: 38.6% and 28.2% of ancestry derived from each subspecies, respectively) and derive the least ancestry from the Sumatran and Malayan populations (mean: 11.1% and 5.0%, respectively; Fig. 1A). Our clustering analyses and PCA results showed that the captive population clusters together by their dominant ancestry in both PC space and the identity-by-state (IBS) tree (Fig. 1B; Supplementary Fig. S6C). We also observe some clustering based on putative geographic origin in the United States and Canada (Fig. 1C), though it is visually less distinguishable than the clustering by ancestry. These results suggest that structure of the Generic population is primarily caused by the breeding events that established the captive population in the United States and there is weak evidence for subsequent geographic structuring, indicating that there is frequent exchange of individuals between breeders. We see further evidence of historical admixture based on mitochondrial haplotype results, where Generic individuals cluster with Amur, Indochinese, and Bengal tiger subspecies. We observed only one Generic sample (SRR836354) grouping with Malayan tigers in mitochondrial haplotype analyses, and none grouping with Sumatran or South China tigers (Supplementary Fig. S2-4). Further information can be found in *Supplementary Notes*.

**Fig. 1:**
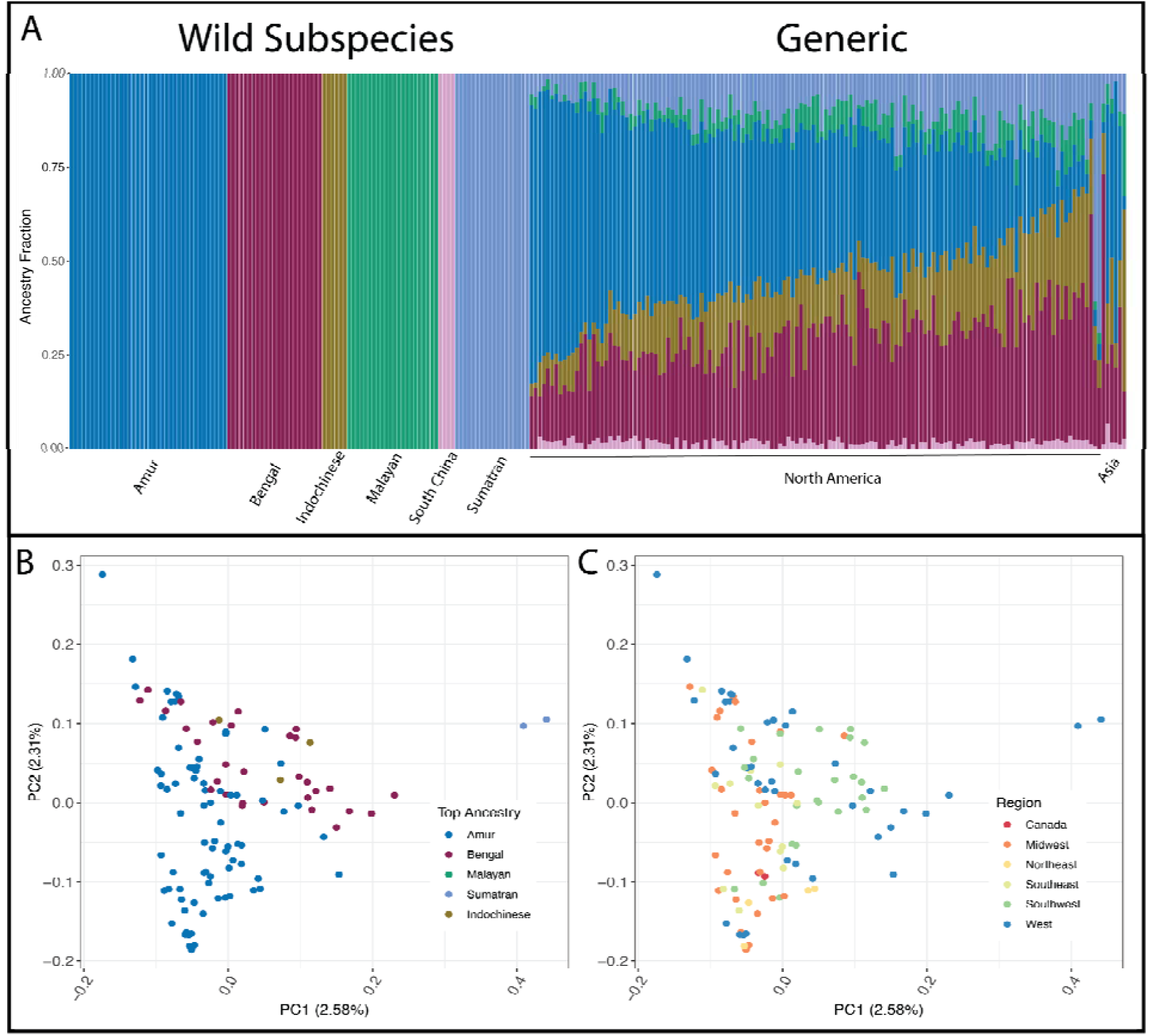
Ancestral diversity in Generic tigers. **A)** ADMIXTURE analysis showing ancestry of all individuals, wild and Generic. **B)** PCA of North American Generic tigers where color corresponds to an individual’s major ancestry component. **C)** PCA of Generic tigers where color corresponds to the region of North America from which the individual originated. For panels B & C one individual was removed as an outlier; details can be found in *Supplementary Notes*.

Interestingly, our analyses also identified several individuals previously labeled as single-ancestry subspecies to be Generic individuals. These individuals were originally labeled as Amur (SRR7651464-67, SRR7651470) and Bengal (SRR836354) subspecies in public repositories, but we observe that they all have a mixed ancestry background based on both PCA (Supplementary Fig. S17) and subsequent admixture analysis (Fig. 1A). These individuals lack additional metadata but are presumed to have originated in Asia (Supplementary Table 1) and thus are labeled as ‘Asia’ in the admixture plot (Fig 1A).

### Tiger subspecies diversity and population history

Since the Generic tigers form a well-admixed population, we next investigated their genetic diversity, inbreeding, and mutation load relative to the single-ancestry, wild subspecies. Primarily, we wanted to determine whether Generic tigers were potential reservoirs of genetic diversity that is absent in the wild, as has been previously suggested. Given that bottlenecks have occurred both in wild populations and during the process of founding the various captive tiger populations, the surviving diversity in the Generic population in comparison to wild populations is unknown. To address this, we estimated the number of distinct alleles per-locus, calculated the proportion of sites that are variant and segregating in the Generic tigers, but fixed in the wild subspecies, and calculated observed heterozygosity for each individual using only unimputed individuals (Fig. 2 & Supplementary Figs. S7-8). ADZE (36) estimates allelic and shared diversity through a rarefaction approach. We show that trajectories for allelic diversity suggested the Indochinese subspecies (Supplementary Fig. S7) contained the most diversity, followed by the Bengal subspecies. However, due to the small sample size for Indochinese tigers (N = 6), it is unclear whether their allelic diversity scales with sample size as it does in the Bengal tiger subspecies (Supplementary Fig. S8).

**Fig. 2:**
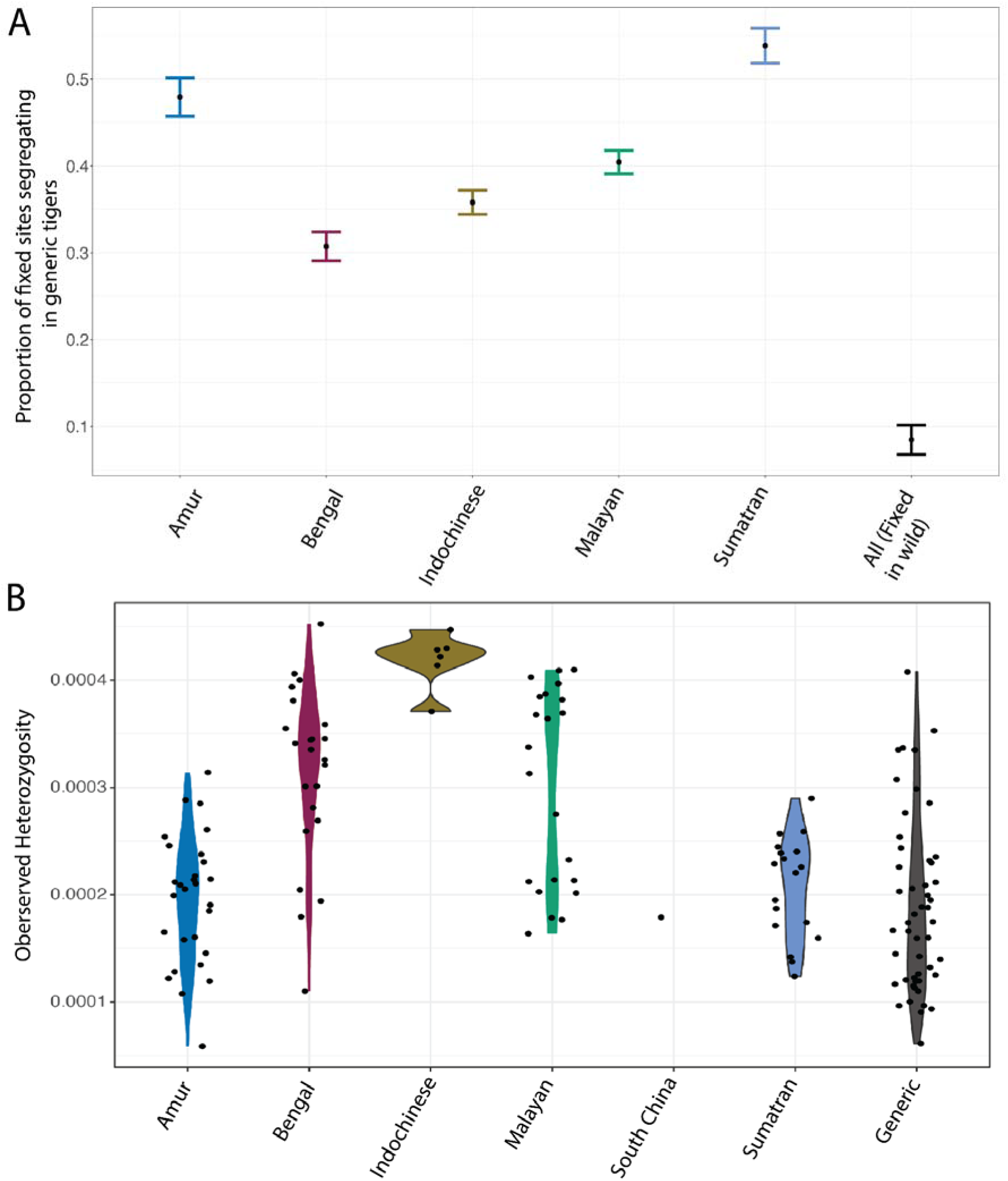
Measurements of shared and within subspecies genetic diversity across tigers. **A)** Proportion of segregating sites that are fixed in each wild subspecies compared to Generic tigers. The mean and standard deviation are computed across 10 replicates of captive individuals (unimputed individuals). South China is not included since there is only one individual. The designation of “All” indicates the site is fixed across all wild subspecies but segregating in the Generic population. **B)** Observed heterozygosity distributions across unimputed individuals, we have applied a horizontal jitter to each data point (See Supplementary Methods and Imputation and filtering for details).

Next, we asked whether there might be an enrichment of variant sites that are solely segregating within the Generic population but fixed within any reference wild subspecies (Fig. 2A). We found that most sites that were segregating within the Generic tigers are also segregating in at least one of the wild subspecies (Fig. 2A). Specifically, less than 10% of sites are exclusively segregating within the Generic tigers (Fig. 2A). Finally, we examined heterozygosity. Generic tiger heterozygosity fell well within the range of other wild subspecies (Fig. 2B). Similar to what we observed with ADZE, Bengal and Indochinese tigers had the highest heterozygosity. Taken together, our analyses suggest that the Generic population is not a major reserve of unique genetic diversity, rather, it contains diversity that is present in the wild.

Due to captive breeding practices and a small founder population, we examined whether the Generic tiger population showed signs of inbreeding. Specifically, we estimated inbreeding using multiple metrics, including runs of homozygosity (ROH), shared identity-by-descent (IBD) segments, and inbreeding coefficients from both SNPs (F_SNP_) and ROH (F_ROH_) using only unimputed data. We observed that the Amur population had, on average, the most Type C (i.e., long runs, informative about recent inbreeding) ROH and highest IBD-sharing (Fig. 3 and Supplementary Figs. S9-10), suggesting that Amur tigers experienced more severe inbreeding than the Generic population. Further, Amur tigers also had the largest amount of Type B (i.e., intermediate length) ROH, which indicates a long-term, small population size (37). Indeed, wild Amur tigers were documented as having experienced an extreme bottleneck in the 1930s/1940s (11, 38), possibly explaining these extreme patterns compared to other subspecies.

**Fig. 3:**
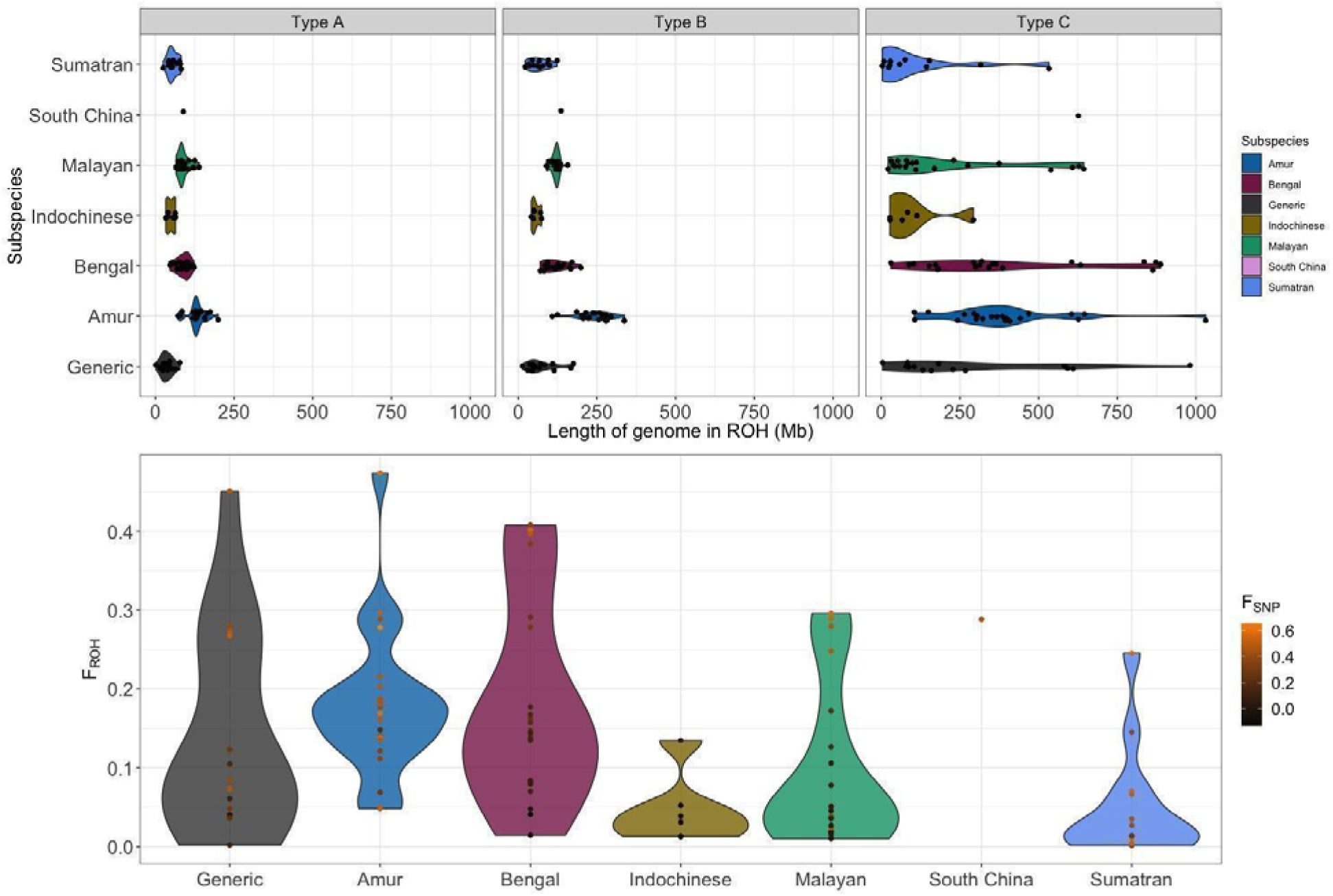
Quantification of different types of ROH across tiger subspecies and measurements of inbreeding. **(Top)** Length of different classes of ROH (A; short, B; intermediate, C; long) for each tiger subspecies and Generic tigers. **(Bottom)** Total proportion of genome in Type C ROH (F_ROH_) for each individual per population. We have labeled each individual with their corresponding inbreeding coefficient measured as F_SNP_, the darkest color (black) is the lowest value, and the brightest color (orange) corresponds to the largest value.

When examining two other metrics of inbreeding (F_SNP_ and F_ROH_) we observed that the Generic tiger population was once again not an outlier. Instead, most Generic individuals had similar inbreeding values to individuals in the other subspecies (Fig. 3). However, there are a few individuals within the Generic population that are quite inbred. The fluctuating levels of inbreeding matched the previous analysis of heterozygosity, where we observed variable levels of genome-wide heterozygosity across the Generic population (Fig. 2B). Additionally, we also found that when examining the site frequency spectra (SFS), the Generic tigers also had the largest proportion of singletons (Supplementary Fig. S11). The large variance in inbreeding coefficients (Type C ROH, F_ROH_, and F_SNP_) suggest that in the case of Generic tigers, admixture between distinct subspecies likely increased heterozygosity in some individuals, but inbreeding within individual facilities eroded it (39). Taken together, our results show that the Generic population is not any more inbred than wild tiger populations, nor is it more diverse.

### Tiger subspecies mutation load

We last sought to characterize the prevalence of putatively deleterious variation in the captive population, relative to each subspecies (unimputed samples). Variants were categorized as putatively deleterious if they were nonsynonymous, derived mutations with a SIFT score less than 0.05 (see *Supplementary Methods* for more detail). We implemented multiple approaches to count deleterious variants in the genome: 1) counting homozygous derived genotypes; 2) counting the total number of homozygous and heterozygous derived genotypes (counting variants); and 3) counting alleles where we tabulate twice the number of homozygous derived genotypes plus heterozygous genotypes (counting alleles).

Given that the Generic population primarily derived its ancestry from the Amur and Bengal subspecies (Fig. 1A), we expected that the distribution of deleterious variation in Generic tigers would be comparable to the Bengal and Amur subspecies. Indeed, the observed distribution of both neutral and deleterious variants in Generic tigers was more similar to the Amur tigers and Bengal tigers than other wild subspecies (Fig. 4 & Supplementary Fig. S12). Overall, we observed similar levels of deleterious variation across Sumatran, Malayan, Indochinese, and Bengal subspecies. Though the mean number of derived deleterious variants was slightly, but consistently, higher in the Sumatran subspecies for all counting methods (Supplementary Fig. S12). We found that, irrespective of subspecies, individuals with the most putatively deleterious derived homozygotes also tended to have the largest inbreeding coefficients, quantified with either F_SNP_ or F_ROH_ (Supplementary Fig. S14). Thus, inbreeding likely led to an overall decrease in fitness for all tigers, irrespective of their ancestry. Our observation that the Generic tigers had the lowest average numbers of derived deleterious mutations, suggests that the Generic tigers may have been able to either purge some of the variants that are deleterious in homozygous form being carried by the other subspecies, or the admixture process reduced their frequency as homozygotes.

**Fig. 4:**
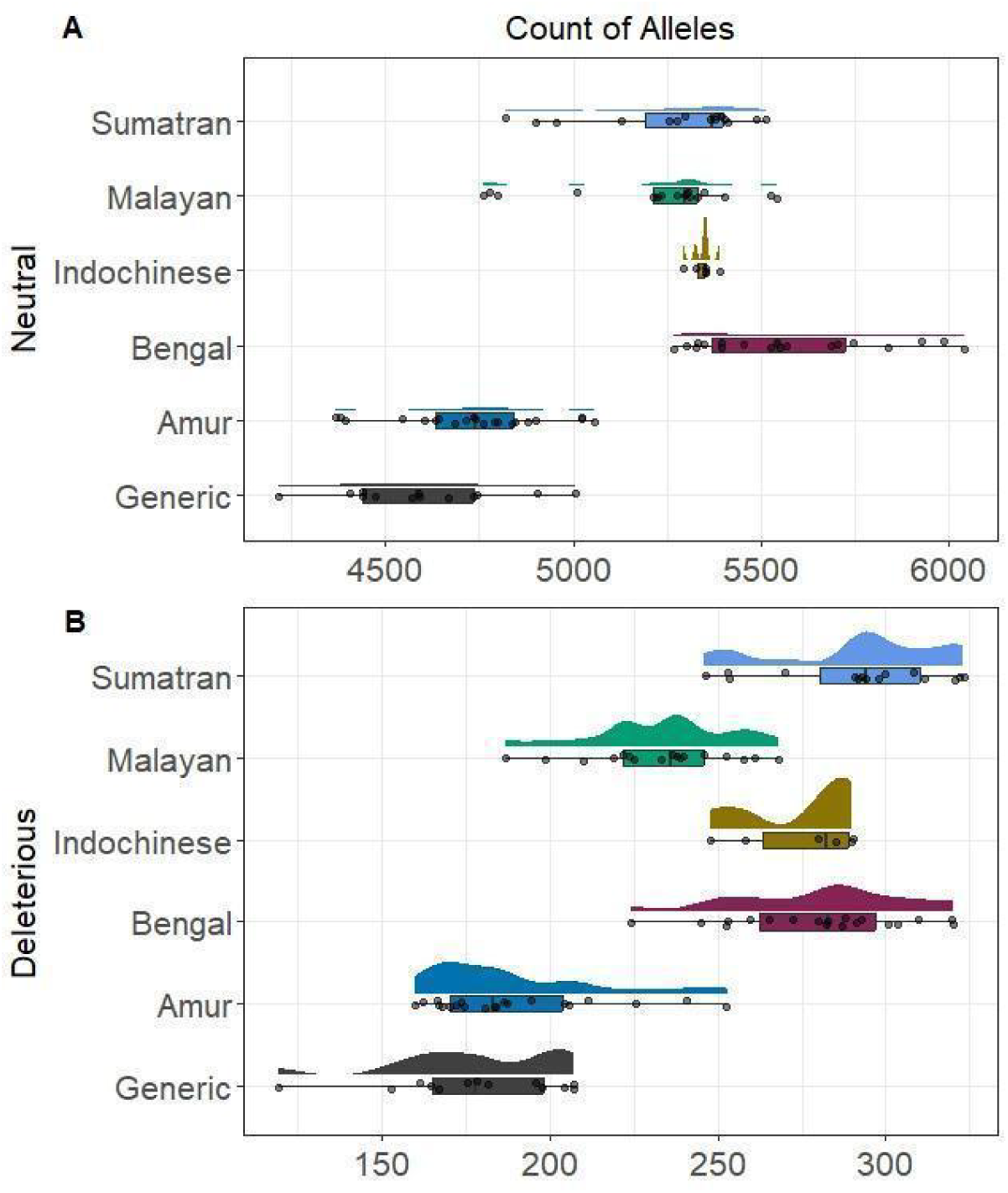
Quantifying genetic load in each subspecies by counting derived neutral and deleterious variants. **A)**: Scaled counts of derived neutral alleles in each tiger subspecies and Generic tigers. **B)**: Scaled counts of derived deleterious alleles in each tiger subspecies and Generic tigers. Extended figure (Supplementary Fig. S12) contains the count of homozygotes and count of variants.

Next, we examined whether putatively deleterious and nonsynonymous homozygous variation tended to be enriched within ROH versus outside of ROH. Recall that counting homozygotes is relevant when we are interested in variants that could be recessive, here we are asking whether putatively deleterious and nonsynonymous variants tend to be in ROH, which would allow them to act recessively and expose them to selection. We identified a significant enrichment of both nonsynonymous and putatively deleterious derived homozygotes within ROH versus outside of ROH across all subspecies (Supplementary Fig. S15). We saw that the Generic tigers were a clear standout in terms of carrying nonsynonymous derived homozygotes within ROH (odds ratio ∼2.9). However, the odds of putatively deleterious derived homozygotes falling within ROH in the Generic tigers are comparable to any of the wild subspecies. In the case of putatively deleterious derived homozygotes within ROH, the standout group became the Indochinese tigers, though they also had the largest 95% CIs (Supplementary Fig. S15). Taken together, these results support that we cannot predict how admixture will affect the burden or distribution of nonsynonymous and putatively deleterious derived homozygotes.

### Creation of a reference panel for future tiger conservation

Given that we collated the largest genomic dataset of tigers to date, we also tested whether the data were sufficient to create an accurate reference panel for low-coverage sequencing and imputation. An imputation pipeline can be particularly useful in wildlife forensics where the ancestry or identity of the individual is unknown, but also for cost-effective sequencing of many individuals. Imputation is generally used in the context of low-coverage data to increase the reliability of genotype calls and improve the accuracy of downstream analyses, and has recently been advocated as an alternative to reduced representation approaches in non-model species, which often amplify a small and potentially biased portion of the genome (40). The primary goal of the reference panel was to assess whether we could accurately impute ultra-low coverage data for the purpose of individual identification and ancestry determination in tigers. The full data was split into a training set (reference panel) composed of 106 single-ancestry individuals (5, 6) with coverage between 2×-43×. Then, imputation was performed using *lowimpute* on 86 low- and ultra-low coverage (0.25×-6×) samples (Supplementary Fig. S1). We tested accuracy in a set of nine individuals (six unimputed and three imputed), where four of the individuals were also represented in the reference panel (See Supplementary Notes for additional details). To test all samples, we randomly downsampled reads from each individual to five different coverages (0.25×, 0.5×, 1×, 2×, 5×) in addition to no down sampling (using all the reads in a standard variant calling pipeline without imputation). Downsampled reads were run through the imputation pipeline, and accuracy of the imputation was tested using variant calls (via non-reference discordance; NDR), ancestry predictions, and the ability to uniquely identify individuals.

We found that as sequencing depth increased, the accuracy of variant calls, as measured by NDR, also increased (Supplementary Table 2). NDR ranged from 7.71% at the highest depth (6×) to 14.49% at the lowest depth (0.25×). We also observed variance in NDR depending on whether a sample was included in the reference panel, as expected, samples from the reference panel had the lowest NDR (Supplementary Table 2). Despite the variance in NDR, ancestry (Supplementary Fig. S16) and the ability to identify individuals (Supplementary Table 4), even at ultra-low coverage (0.25×), remained accurate. Our results show that low-coverage sequencing and imputation are a sufficient and low-cost alternative to high-coverage sequencing or genotyping for the purposes of ancestry and individual identification using this reference panel for tigers.

## Discussion

### Admixture and variation

We found that all Generic tigers sampled from the U.S. are admixed and contain ancestry from all six extant tiger subspecies (Fig. 1A). Most Generic tigers derive a majority of their ancestry from the Amur and Bengal subspecies (Fig. 1A), but individuals varied in the proportion of ancestry derived from any one subspecies (Fig. 1A, Supplementary Fig. S5). The length distribution of local ancestry segments (Supplementary Fig. S5) indicated that there are likely very few individuals in the extant U.S. Generic tiger population that have directly descended from recently introduced, single-ancestry sources. The timing and sources of these introductions is unknown. Though we observe ancestry from South China tigers in many of the captive samples, the low number of available samples for South China generally supports additional investigations before definitive conclusions can be made about their ancestry contributions to the Generic population.

Our findings differ from the results obtained from Luo et al. 2008, where 30 microsatellite loci and mitochondrial (4kb) data were used to investigate samples from captive tigers (N = 105) obtained between 1982 and 2002 from 14 countries (including the U.S.). By comparing their data to voucher specimens (39 Amur; 2 South China; 33 Indochinese; 22 Malayan; 17 Bengal; 21 Sumatran), they found that 49 of the captive tigers belonged to single subspecies ancestry, while 52 were found to be of admixed ancestry (24). A total of 18 tigers did not match the suspected ancestry (i.e. the tiger was found to be admixed compared to the source reporting single subspecies ancestry or vice versa). Luo and colleagues (2008) did not describe the origin of the sampled individuals in detail, but it is possible that individuals of single-subspecies ancestry were being actively mixed into the privately-owned captive tiger population when those samples were collected. Alternatively, it is possible that the small number of microsatellites used in the previous study have comparatively limited power to distinguish the complex ancestry of the Generic tigers compared to thousands of SNPs, as has been the case in other systems (41–43). Interestingly, Luo and colleagues (2008) found that captive tigers with mixed subspecies ancestry primarily contained ancestry from the Indochinese tiger lineage, whereas the tigers analyzed herein contained mostly Amur and Bengal ancestry. Additional genomes for captive non-zoo individuals from different countries would provide insight into whether tiger ancestry proportions are a useful fingerprint distinguishing continental or country-specific tiger populations that could be leveraged for wildlife investigations.

While there is no doubt that the Generic tiger population carries some unique variation simply because of the large number of individuals in captivity (i.e., census size), we have shown in our sampling that the population does not contain more variation or more unique genetic variation than its single-ancestry counterparts (Fig. 2, Supplementary Figs. S7-8), contrary to previous studies (24). In other words, Generic tigers have average heterozygosity (Fig. 2B), possess very few unique segregating sites (Supplementary Fig. S7-8), and when compared to other subspecies, most sites that are segregating in the Generic population are also segregating in the wild populations (Fig. 2A ). While there may be some beneficial alleles circulating in the Generic population, predicting which loci are adaptive is a challenging task for even the most well-studied species, and implementing gene-based management strategies can also bring about harmful side-effects (44, 45).

### Deleterious variation

The Generic population contains less deleterious recessive homozygous variation and relatively low amounts of Class C ROH relative to wild populations (Fig. 3 and Fig. 4). Given the degree of admixture in the Generics, the low frequency of deleterious mutations is expected, yet surprising given the purported inbreeding of ‘farmed’ tigers (46). However, Generic tigers also have the largest enrichment of nonsynonymous variation (counting alleles, counting variants, counting derived homozygotes) within ROH versus outside. In other words, though Generic tigers have a comparable amount of nonsynonymous and putatively derived homozygotes compared to their wild ancestors, they tend to carry more of the nonsynonymous variation within ROH than their ancestors. Thus, the results of admixture on the genomic landscape in Generic tigers are, in this case, not obvious. Heterozygosity and load are similar across all tigers, likely due to a composite effect of admixture between distinct lineages increasing heterozygosity, followed by severe bottlenecks of repeated founding events and deliberate inbreeding. The observed outcome of admixture in the Generics is the result of chance but is not necessarily predictive of other metrics of inbreeding, load, or heterozygosity, and also does not negate possible effects of outbreeding, which we did not examine here. The increase in nonsynonymous load within versus outside of ROH, and population-level heterozygosity falling between major ancestry sources has also been observed in human populations, specifically in admixed Latin American founder populations (39).

### Ancestry inference for unknown individuals

The final objective of this study was to compile a reference dataset and imputation panel capable of identifying the ancestry and identity of individuals. At present, tiger forensic work relies primarily on mitochondrial and microsatellite data (47). Mitochondrial markers in felid lineages contain nuclear mitochondrial inserts (numts), making interrogating these regions difficult and error-prone (48, 49). Further, mitochondrial markers are only indicative of the maternal lineage of an individual and are not able to identify individuals with mixed ancestry. For complex admixture scenarios, examining only a handful of microsatellites may cause biased results or in the case of species or subspecies-level hybridization distort ancestry signals (50–53). As sequencing costs decline, arrays and panels are set to emerge as the predominant and cost-effective way to query forensic and low-quality samples, compared to microsatellites (54, 55). Arrays and panels require either a sufficient number of SNPs (56, 57) or if a reduced set is needed to be cost-effective, a very careful selection of SNPs which are reflective of ancestry or relatedness. Low-coverage WGS provides an avenue for rapid and simple identification, which could be performed on any number of sequencing machines from Illumina to the portable Oxford Nanopore. The reference dataset and imputation pipeline from our study provide a robust way to identify individuals and their ancestry rapidly and cost-effectively, even from very low-coverage samples.

## Conclusions

There has been ongoing discussion regarding how captive populations might contribute to conservation, and particularly whether they harbor unique alleles beneficial to the survival of the species or represent historical variation that is no longer present in the wild. Here, we have shown that the captive, privately-owned tiger population does not contain significant unique variation compared to the wild subspecies. Additional investigations into the remaining diversity in captive zoo populations are unlikely to reveal additional diversity since many of these lineages are the putative founders of the Generic tiger population. Several notable zoo populations, such as giraffes (58) and orangutans (59) have sourced animals for captive breeding programs from multiple, genetically distinct populations, subspecies, or species. If we continue to consider captive populations as diversity reservoirs for wild species, additional investigation into the potential outcomes and interplay of such processes (admixture, inbreeding vs outbreeding depression) is extremely important. We further caution that deleterious variation analyses, especially in non-model organisms, should be viewed skeptically. Deleterious variation analyses generally rely on annotations from other organisms and the accurate inference of an ancestral state, both of which can be imprecise analyses even in the most well-studied model organisms (60–62). Computational annotations take a holistic approach to determining the likely consequences of variation, but until functional validation of variants occurs, genetic load should be integrated into management plans with extreme caution.

Cumulatively, our analyses are concordant with the known history of tiger subspecies in terms of historic bottlenecks and recent inbreeding, yet we also reveal the unique history of the large population of Generic tigers in the United States and provide a comprehensive overview of all extant subspecies and available data. Most Generic tigers contain ancestry from all six wild tiger subspecies in their genomes. Contrary to previous hypotheses, most Generic tigers do not show signs of severe, recent inbreeding, nor do they hold unique diversity. Thus, the role they might play (if any) in terms of conservation of genetic diversity is unclear. Barring outbreeding concerns, we find no genetic reason why Generic tigers from the U.S. would not be useful in augmenting tiger populations, since most individuals do not contain an excess of deleterious mutations or appear to be inbred. Whether it is wise to keep the tiger subspecies separate to preserve their genetic uniqueness, or whether certain circumstances warrant lineage mixing remains to be seen for the tiger and many other species. South China tigers have already been genetically rescued by another subspecies (63), and what many populations may require for survival is more individuals, but where they are sourced is a critical choice. For example, here we showed that the Sumatran tiger population in particular may be in need of conservation action due to an excess of homozygous deleterious alleles (Fig. 4 Supplementary Fig. S12 & 13), but further investigation into the single-ancestry captive populations will further elucidate the genetic resources available for tiger conservation and simulations could help determine the best sources for possible translocations (33).

The U.S. Generic population is the product of unique evolutionary admixture events which may help us better understand the consequences of mixing divergent, wild lineages in the future. To encourage future study and aid illegal trade and trafficking investigations, we also present a reference panel that in conjunction with imputation can accurately identify the ancestry and identity of an individual using even ultra low-coverage sequencing, demonstrating the validity of this approach. Low-coverage WGS is an attractive alternative to creating marker panels, which often are affordable and accessible only with caveats (e.g. when running many markers or many samples), that often exclude conservation budgets and sample needs. Taken together, our results are not only informative for the conservation management of tigers, but also provide a cost-effective path for others in the conservation community to follow for generating a reference panel when using low-coverage sequencing for identification of individuals and their ancestral origins.

## Methods

### Sample collection and sequencing

A total of 154 tiger samples were collected opportunistically during routine vet care from sanctuary facilities by vet and sanctuary staff or from existing biobank collections (Supplementary Table 1). All samples were extracted using a Qiagen DNeasy kit (Cat. No. 69504) and samples prepared using a modified Nextera library prep protocol (64). 77 of these samples (listed as ‘Unimputed’ in Supplementary Data 1) were sequenced between approximately 2× and 5× depth. The remaining 77 samples (‘Imputed’ in Supplementary Data 1) were sequenced at approximately 0.25×. Tigers collected in sanctuaries or with unknown ancestry are labeled as “Generic” for the purposes of this manuscript. We additionally collected data on the putative location of birth of each tiger from the sanctuaries in North America. These locations were then translated into one of six regions: West (Washington, Oregon, California, Nevada, Montana, Idaho, Wyoming, Colorado, Utah), Southwest (Arizona, New Mexico, Texas, Oklahoma), Midwest (North Dakota, South Dakota, Nebraska, Kansas, Minnesota, Iowa, Missouri, Wisconsin, Illinois, Michigan, Indiana, Ohio), Southeast (Arkansas, Louisiana, Tennessee, Mississippi, Alabama, Georgia, Florida, Kentucky, West Virginia, Virginia, North Carolina, South Carolina), and Northeast (Maine, New Hampshire, Massachusetts, New York, Pennsylvania, Maryland, Delaware, Connecticut, Vermont, Rhode Island, New Jersey) and Canada (Canada was not subdivided since N = 2 and both individuals were from the same facility).

### Variant-calling and reference panel construction

An additional 100, publicly available (as of December 2019) tiger genome samples were downloaded from NCBI. Reads were mapped to the GenTig1.0 genome (65) using BWA-MEM v0.7.17 (66) and variant calling was subsequently performed by Gencove using the Genome Analysis Toolkit (GATK) v4.1.4.1 (67) according to best practices. Initial variant calling was performed on all samples available at the time, excluding those sequenced at 0.25×. All of these samples are referred to as ‘unimputed’ for the purposes of this manuscript. Initial variant calling yielded a total of 23,579,569 variable sites across the entire genome. We restricted calls to biallelic sites using BCFtools v1.6 (68), and subsequently filtered for quality, missingness, and depth. See *Supplementary Methods* for additional details. The final data set contained a total of 7,519,430 sites across unimputed individuals and a callable genome of 2,174,711,735 base pairs.

In order to select individuals to build the reference panel and accurately split individuals into groups for kinship estimation, we conducted Principal Component Analysis (PCA) to ensure that all individuals in the unimputed dataset were clustering according to subspecies using PLINK v2 (69). Individuals were subsequently split into ancestry groups to form the reference panel, which included representatives from all six tiger subspecies, but no Generic individuals. Further, we tested several methods for detecting relatedness using pedigreed individuals in the dataset, which were subsequently used to identify and remove duplicates. Additional information can be found in *Supplementary Methods*.

### Imputation and filtering

Using only single-subspecies ancestry individuals verified here and by previous studies, we developed a reference panel to impute variants for an additional 86 individuals (labeled as ‘imputed’ in Supplementary Data 1) through the *loimpute* pipeline developed by Gencove and available at www.gencove.com (70). The 86 imputed individuals were composed primarily of individuals sequenced at ultra-low coverage (N = 75; 0.25×), but also included two individuals from a Canadian Zoo sequenced at approximately 3×, and an additional 9 samples that became publicly available after the initial variant calling had been performed (see Supplementary Data 1 for details).

We combined the VCFs for imputed individuals with the unimputed individuals using BCFtools *merge*. Because imputation emits a call for every site in the reference pipeline, we restricted the merged sites VCF to retain only the quality sites identified after initial variant calling and filtering. We further checked for imputation accuracy using concordance measures, specifically non-reference discordance (71), and examined the accuracy of ancestry and relatedness measures over a variety of coverages. Additional information can be found in *Supplementary Methods*.

### Genetic diversity

We used several approaches to investigate genetic diversity across tigers. The first approach was to create equal sized groups of (N=10) individuals and tabulate the proportion of sites that were SNPs in the Generic population and fixed in each wild subspecies. We used the same reference groups from each subspecies and generated 10 replicate samples (with replacement) of the Generic tigers to check whether these proportions varied across the Generic population.

Observed heterozygosity was also calculated as the total number of heterozygous sites divided by the number of callable sites (2,174,711,735) in the genome. Observed homozygous sites were counted in each subspecies using VCFtools (72) using the ‘--het’ flag and heterozygous sites were calculated by subtracting the (O)HOM (observed homozygosity) column from the NSITES (the total sites queried) column. The number of callable sites was determined as the total number of base pairs minus the sites with mappability scores < 1 for autosomal scaffolds. We additionally tested to see if heterozygosity was correlated with missingness. See *Supplementary Methods* for details.

### Runs of homozygosity (ROH)

GARLIC(73) was used to detect ROH. The error was set at 0.001, the window size at 700, and centromeres were set as 0,0 since no centromere information was available. To ensure that ROH were not mistakenly called on regions with an excess of missing calls, each ROH was then intersected with the number of callable sites in each window (see *Heterozygosity* section for details). ROH larger than 100kb and containing callable sites within one standard deviation (±0.066) of the mean coverage (0.655) were retained. ROH were binned into different size classes A (short), B (intermediate), and C (long) per subspecies. Binning was based on the use of a Gaussian mixture function that fits a model to the ROH length distribution within the group. Type A ROH are typically indicative of linkage-disequilibrium blocks. Type B ROH are informative about long-term small population sizes and cryptic relatedness. The presence of Type C ROH indicates recent inbreeding in the population. F_ROH_ was computed as the total fraction of the genome within a type C ROH.

### Identity-by-descent segments (IBD)

IBD segments were called using TRUFFLEv1.38 with parameters ‘--segments --missing 1 --maf 0 --nofiltering’ (74). Segments greater than 2Mb where the fraction of coverage by callable sites (count listed above) was within one standard deviation (±0.032) of the mean coverage (0.660) were retained. IBD scores were computed using the same approach from Nakatsuka et al. (75), where we normalize shared IBD by the number of sampled individuals.

### Putatively neutral and deleterious variation

We polarized and annotated sites by first filtering the data to scaffolds that corresponded to main chromosomes from felCat8 (76); GCF_000181335.2). Coordinates were identified using liftOver (77). The remaining sites, with felCat8 coordinates, were annotated with an impact and consequence using VEP v92. We removed intergenic sites, splice acceptor, splice donor, splice region annotations, and selected the most damaging impact for a given transcript. Sites were coded as nonsynonymous (NS), synonymous (SYN), or loss of function (LOF) and SIFT scores were added (78). Information from VEP was combined with a SIFT score to find putatively neutral (SYN with SIFT score greater than 0.05) and putatively deleterious sites (NS or LOF with SIFT score less than 0.05). See *Supplementary Methods* for details about annotations and SIFT scores.

Ancestral bases were identified with Progressive CACTUS (79) alignment of the GenTig1.0 genome with 9 other felid genomes; further details can be found in *Supplementary Methods*. To assess load in each subspecies, we used only unimputed individuals with at least 5× coverage and less than 5% missing data. Lastly, we scaled the number of sites per individual by subtracting the total number of variant sites across all individuals from missing sites to get the total number of called sites. Each count was divided by the number of callable sites for that individual; the proportion was multiplied by the average number of callable sites across all subspecies.

## Data Availability

Whole genome sequence data associated with this study has been deposited into NCBI under bioproject number PRJNA976043 and will be released upon acceptance. Raw variant call files (VCFs) have been uploaded to dryad repository https://doi.org/10.5061/dryad.k0p2ngff1.

## Code Availability

All code associated with this project can be found at https://github.com/jaam92/Tigers/.

## Acknowledgements

We would like to acknowledge each of the sanctuaries and zoos who contributed samples, knowledge, and time to this study, specifically the Performing Animal Welfare Society, the Exotic Feline Rescue Center, In-Sync Exotics, Turpentine Creek Wildlife Refuge, Carolina Tiger, and others. We especially acknowledge Joe Taft and Rebecca Rizzo of EFRC, Jackie Gai and Ed Stewart of PAWS, and Vicky Keahy of In-Sync, who have spent many hours discussing this project with us. This project would not have been possible without Bill Nimmo and Kizmin Reeves of Tigers in America. In addition, we thank the San Diego Frozen Zoo for access to their collections and their collaborative spirit. We also thank Sara Huston Katsanis for procuring the Carolina Tiger Rescue and Tiger World specimens and supporting the mentoring grant for V. Grant.

## Author contributions

EEA, EAH, and UR conceived the study; KP, TY, VBG, and EEA generated the sequence data; EEA, JM, GG, KAS, BYK, CBK, JKP analyzed the data. All authors contributed to editing, guiding analyses, and writing the manuscript.

