## Supplementary Information for "Unraveling the Genomic Diversity and Admixture History of Captive Tigers in the United States"

### Supplementary Methods

#### Variant calling and reference panel construction

At the time of the reference panel construction, an additional 99 tiger genome samples were publicly available and downloaded from NCBI. Reads were mapped to the GenTig1.0 genome^1^ using BWA-MEM v0.7.17^2^ and variant calling was subsequently performed by Gencove using the Genome Analysis Toolkit (GATK) v4.1.4.1^3^ according to best practices. Initial variant calling was performed on all samples, excluding those sequenced at 0.25×, for a total of 177 individuals.

We filtered for low-quality sites using BCFtools v1.16^4^. We first restricted to biallelic sites using BCFtools *view*, with the flags ‘-m2, -M2, -v SNPs’. We next examined summary statistics (depth, allelic number) of the data using BCFtools *query* -f. To exclude low-quality sites, we filtered first based on site missingness, using AN > 317. We then restricted the data to only autosomes, and further filtered based on depth and quality using BCFtools *view* and included sites that had a quality score of at least 20, a minimum depth across individuals of 177× and a maximum depth across individuals of 3000× (the average depth across all individuals per biallelic SNP site was 1725×). We last used a mappability filter to remove sites with low mappability. We estimated mappability scores using Genmap v1.3.0^5^. The reference file was first indexed using genmap *index* -F, and the mappability subsequently indexed using genmap *map* with flags ‘-K 30’, ‘-E 2’, and ‘-b’. The resulting bedgraph file of mappability scores was then filtered to exclude sites that had a mappability score <1 using filterGM.rb v0.3.2 (from RatesTools^6^, https://github.com/campanam/RatesTools/). These sites were then filtered from the VCF using VCFtools v0.1.15^7^  ‘--exclude-bed’.

In order to select individuals to build the reference panel and accurately split individuals into groups for kinship estimation, we conducted Principal Component Analysis (PCA) to ensure that all individuals in the unimputed dataset were clustering according to subspecies using PLINK v2^8^ with flags ‘--bfile’, ‘--allow-extra-chr’, and ‘--pca 10’. Individuals were subsequently split into ancestry groups to form the reference panel, which included representatives from all six tiger subspecies. Further, we tested several methods for detecting relatedness using pedigreed individuals in the dataset, which were subsequently used to identify and remove duplicates. Additional information can be found in Supplementary Methods, Relatedness.

#### Imputation and filtering

Using only the putative single subspecies ancestry individuals verified above, we developed a reference panel to impute variants for an additional 86 individuals (labeled as ‘imputed’ in Supplementary Data 1) through the *loimpute* pipeline developed by Gencove and available at [www.gencove.com](http://www.gencove.com)^9^. The pipeline is based upon algorithms based on the copying model of Li and Stephens^10^. The 86 imputed individuals were composed primarily of individuals sequenced at ultra low-coverage (N = 75; 0.25×), but also included two individuals from a Canadian Zoo sequenced at ~3×, and an additional 9 samples that became publicly available after the initial variant calling had been performed (see Supplementary Data 1 for details). Because the Gencove imputation pipeline sets a maximum depth it will allow (6×), these 9 individuals were downsampled prior to being imputed using the seqtk v1.322 (https://github.com/lh3/seqtk) pipeline with the command seqtk *sample* ‘-s100’. We aimed for a depth of approximately 5×. The final average coverage for these individuals can be found in Supplementary Data 1.

We combined the files for imputed individuals with the unimputed individuals using BCFtools v1.16^4^ *merge*. Because imputation emits a call for every site in the reference pipeline, we restricted the merged sites VCF to retain only the quality sites identified after initial variant calling and filtering using BCFtools *view* with flag ‘-R’. We further checked for imputation accuracy using concordance measures and examined the accuracy of ancestry and relatedness measures over a variety of coverages. Additional information can be found in *Supplementary Notes*.

#### Mitochondrial haplotypes

Whole genome sequence data was mapped to a tiger mitochondrial reference genome (MH124106.1) using BWA-MEM v0.7.17^2^ and sorted using Samtools v1.8^11^. Reads that mapped to the mitochondrial reference genome were extracted and converted to paired FASTQ files using the *bamtofastq* function in BEDTools v2.27.1^12^. To remove reads that were likely from nuclear mitochondrial inserts (numts)^13^, we made a new reference file consisting of the mitochondrial reference genome and a numt reference (DQ151551.1). Paired FASTQ files were mapped to this new reference file, sorted, and consensus sequences generated using ANGSD v0.931^14^.

Consensus sequences for the mitochondrial reference genome were aligned in GeneiousPrime v.2020.1.1 with the published sequence data from Luo et al. 2004^15^ (AY736559–AY736808). Whole mitochondrial genome sequences were trimmed to the 10 gene regions (4,078bp) used in Luo et al. 2004^15^. Any samples with more than four bp of missing data were removed from the alignment. We additionally screened for the presence of numt sequence contamination, by counting the number of SNPs compared to the most common haplotype in each sample in each gene. Samples that displayed more than twice as many SNPs in a gene than observed in any of the reference haplotypes were considered to have numt contamination in that gene and were subsequently removed. Haplotype networks were constructed using Median-joining networks in PopArt^16^. Haplotype networks were also generated after removing only the genes (rather than individuals) with numt contamination to retain more samples in the dataset.

#### Heterozygosity

We created equal sized groups of (N=10) individuals across all subspecies. Then we counted the total number of sites that were SNPs in the generic population and fixed in any wild subspecies. Next, we kept the same reference groups of wild tigers, and generated 10 replicate samples (with replacement) of the generic tigers to check whether these counts varied across individuals in the captive population.

Heterozygosity was then calculated as the total number of heterozygous sites divided by the number of callable sites in the genome. Observed homozygous sites were counted in each subspecies using VCFtools using the ‘--het’ flag and exported into R, and heterozygous sites were calculated by subtracting the (O)HOM column from the NSITES column. The number of callable sites was determined as the total number of base pairs minus the sites with mappability scores < 1 (See *Reference panel construction* section for details) for autosomal scaffolds. We additionally tested to see if heterozygosity was correlated with missingness. Using VCFtools, we calculate the proportion of missing sites per individual using VCFtools ‘--missing-indv’.

#### Relatedness

Relatedness was estimated for all individuals using SNPRelate’s IBDMLE function (see Supplementary Notes) and validated using IBD sharing and pedigrees when available. The unrelated and unimputed individuals with greater than 5× coverage from each subspecies were identified and used for all analyses with ROH and IBD. We consider unrelated individuals as at most 3rd degree relatives.

#### Local ancestry

To investigate local ancestry, we used the set of phased reference files generated for the imputation pipeline (duplicate individuals were removed). To infer local ancestry across all captive individuals, we used the software RFMix v2.03-r0^17^ and assumed a genetic map of 100Mb/1cM. Because RFMix requires multiple individuals, we removed the single South China individual for local ancestry analysis. RFMix was run using default parameters and results plotted using ggplot2 v3.3.6^18^.

#### Runs of homozygosity (ROH)

Only unimputed, unrelated, individuals with greater than 5× coverage were used. We first converted the VCF files to PLINK format using plink v1.9^8^ with the VCF files as input and the ‘--recode’, ‘--const-fid’, and the ‘--allow-extra-chr’ flags. Subsequently, we used the software GARLIC^19^ to detect ROH in each subspecies and the generic tigers. The error was set at 0.001, the window size at 700, and centromeres were set as 0,0 since no centromere information was available.

To ensure that ROH was not mistakenly called on regions with an excess of missing calls, each file was then intersected with a callable sites file (see *Heterozygosity* section for details). Only ROH larger than 100kb and containing callable sites within one standard deviation (0.066) of the mean coverage (0.655) were retained. ROH were divided into different size classes A (short), B (intermediate), and C (long) per subspecies. Binning is based on the use of a Gaussian mixture function that fits a model to the ROH length distribution within the group. Type A ROH are typically indicative of linkage-disequilibrium blocks. Type B ROH are informative about long-term small population sizes and cryptic relatedness. Lastly, the presence of Type C ROH indicates recent inbreeding in the population. F_ROH_ was computed as the total fraction of the genome within a type C ROH. The genome length used was the number of callable sites, which was 2,174,711,735 base pairs.

#### Identity-by-descent segments

Only unimputed, unrelated, individuals with greater than 5× coverage were used. Identity-by-descent (IBD) segments were called using TRUFFLEv1.38^20^ with parameters ‘“--segments --missing 1 --maf 0 –nofiltering’ ^20^. The extra TRUFFLE parameters allow us to convert start and end positions from the output segment file back to positions in the original VCF file. After IBD segments were called, we intersected each segment with the total callable variant and invariant sites, to find the total fraction of the IBD segment that was covered. After converting positions back to the VCF coordinates, only segments greater than 2Mb and where the fraction of coverage by callable sites (count listed above) was within one standard deviation (0.032) of the mean coverage (0.660) were retained. IBD scores were computed for each subspecies and the generics using the same approach from Nakatsuka et al.^21^.

#### Site frequency spectrum (SFS) and polarization

First, all individual felid genomes were extracted from a 241-way mammalian alignment^22^. The *Panthera* *pardus*, *Panthera onca*, *Felis* *catus* (specifically, FelCat8), *Felis* *nigripes*, *Puma* *concolor*, and *Acinonyx* *jubatus* genomes were used as-is. We replaced the PanTig1.0 genome with the more contiguous GenTig1.0 genome. Additionally, we included lion (*P. leo* ^23^), snow leopard (*P.* *uncia* ^24^), and clouded leopard (*Neofelis* *nebulosa*, unpublished, courtesy G. Barsh, C. Kaelin) genomes. The whole-genome alignment was performed with Progressive Cactus^25^ , which takes a guide tree alongside whole genome sequences and reconstructs ancestral genome sequences for each node in the tree during the alignment process. The following cladogram was provided as the guide tree:

(((Panthera_tigris:0.005, (((Panthera_pardus:0.005,Panthera_leo:0.005):0.005,Panthera_onca:0.005):0.005,Panthera_uncia:0.005):0.005):0.005,Neofelis_nebulosa:0.005):0.005,(((Felis_catus:0.005,Felis_nigripes:0.005):0.005,Puma_concolor:0.005):0.005,Acinonyx_jubatus:0.005):0.005).

We leveraged the ancestral genome reconstruction for the common ancestor of the tiger and the (((Panthera_pardus,Panthera_leo),Panthera_onca),Panthera_uncia) clade to polarize each variant call. Therefore, the ancestral base was defined as the base in the common ancestor of tigers and other big cats. Progressive Cactus identifies the ancestral bases on the phylogenetic tree via maximum-likelihood assuming a Jukes-Cantor model of substitution. For sites where the tiger was homozygous, we used the Progressive Cactus allele as the ancestral allele. For sites where the tiger was heterozygous and one of the alleles matched the Progressive Cactus reference allele, we used that allele as the ancestral allele. If neither allele in the VCF matched, we removed the site.

We used only unimputed, unrelated, individuals with greater than 5× coverage to create the SFS. We created two groups, one with N=10 unrelated individuals and second with N=6 unrelated individuals, which are a subset of the N=10 group. We created the two groups to keep the Indochinese population which has a limited sample. Unfortunately, we were forced to drop the South China population since there is only a single unimputed sample.

#### Putatively neutral and deleterious variation

To assess load in each subspecies and in the generic tigers, we used only unimputed individuals with at least 5× coverage and kept individuals with less than 5% missing data. In order to polarize the data and annotate sites, we subset the tiger data to only include scaffolds that corresponded to autosomes from felCat8^26^ (GCF_000181335.2). Coordinates were identified using liftOver^27^. Then we input remaining sites with the felCat8 coordinates into VEP^28^ v92 and annotated each site with an impact (“LOW”, “MEDIUM”, “HIGH”) and consequence. Next, we removed all intergenic sites, splice acceptors, splice donors, splice region annotations, and selected the most damaging impact for a given transcript. We coded each site as nonsynonymous (NS), synonymous (SYN), or loss of function (LOF). We classified the following annotations as loss of function: 
"Stop gained, splice region variant”, "stop lost”, “start lost”, “start lost, synonymous variant", "stop gained, start lost", “stop gained". Next, we added SIFT scores to each variant^29^. SIFT^29^ scores were added by downloading scores from felCat5^26^ and lifting each position over to felCat8^26^ coordinates. Information from VEP was combined with a SIFT score to find putatively neutral (SYN with SIFT score greater than 0.05) and putatively deleterious sites (NS or LOF with SIFT score less than 0.05).

In other words, the total number of sites that were annotated was 50,060 and we retained individuals with less than 2,500 sites annotated as missing. Then, we scaled the number of sites for all individuals. We scaled sites per individual by subtracting the total number of variant sites across all individuals from missing sites to get the total number of called sites. Next, we divided each count by the number of callable sites for that individual. Lastly, we multiplied the proportion by the average number of callable sites across all subspecies.

### Supplementary Notes

#### Population structure and global ancestry

To examine variation across wild and captive tigers we ran a final PCA analysis (N = 255) with duplicates removed and individuals placed in correct ancestry groups (Supplementary Fig. 1). Supplementary Fig. 1 shows the top 3 PCs across these groups, and we can see clear clustering of wild tigers and the captive tigers. We can also see that the captive tigers are dispersed across PC space and that each subspecies forms its own unique cluster.

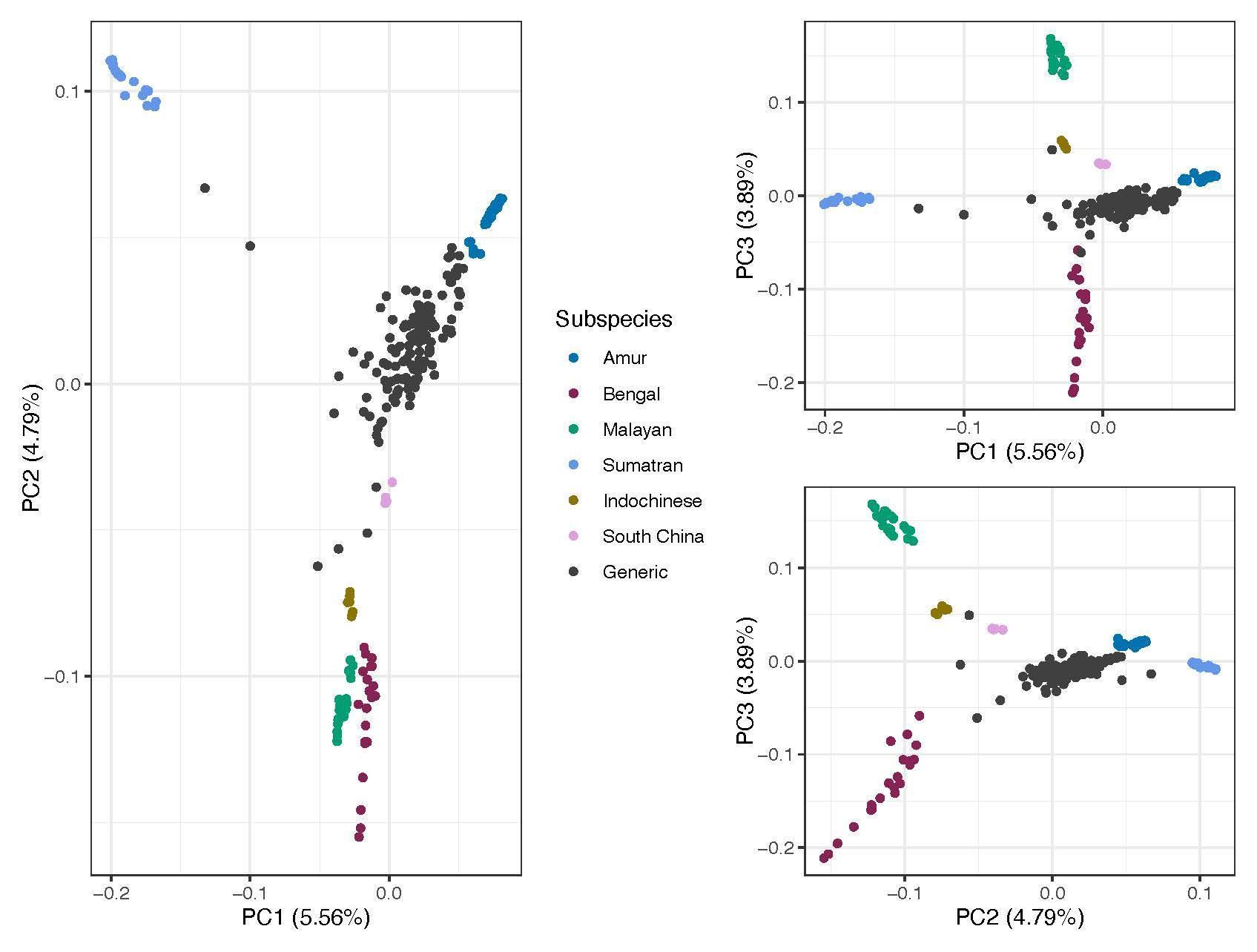

**Supplementary Fig.** **1** Top 3 PCs for final set of individuals, not including duplicates and with misidentified individuals reassigned to the correct population.

#### Mitochondrial Diversity

We generated mitochondrial consensus sequences by mapping to a tiger mitochondrial reference genome (See Supplementary Methods). We restricted our analyses to 10 genes contained in Luo et al. 2004^15^. Contamination was only observed in two genes. Specifically, ten samples had numt contamination in Gene 1 (AMU11, PAWS5, ISE14, EFRC46, AMU4, EFRC1, ISE3, PAWS4, ISE12, EFRC11) and one sample had numt contamination in Gene 2 (ISE13).

A median joining haplotype network was constructed for the dataset after removing all samples with numt contamination in any gene (Supplementary Fig. 2) using PopArt^16^. A haplotype network was also constructed after trimming Gene 1 from the dataset and removing the one sample with numt contamination on Gene 2 (Supplementary Fig. 3). Lastly, we generated a haplotype network after trimming Gene 1 and 2 from the dataset and retaining all samples (Supplementary Fig. 4).

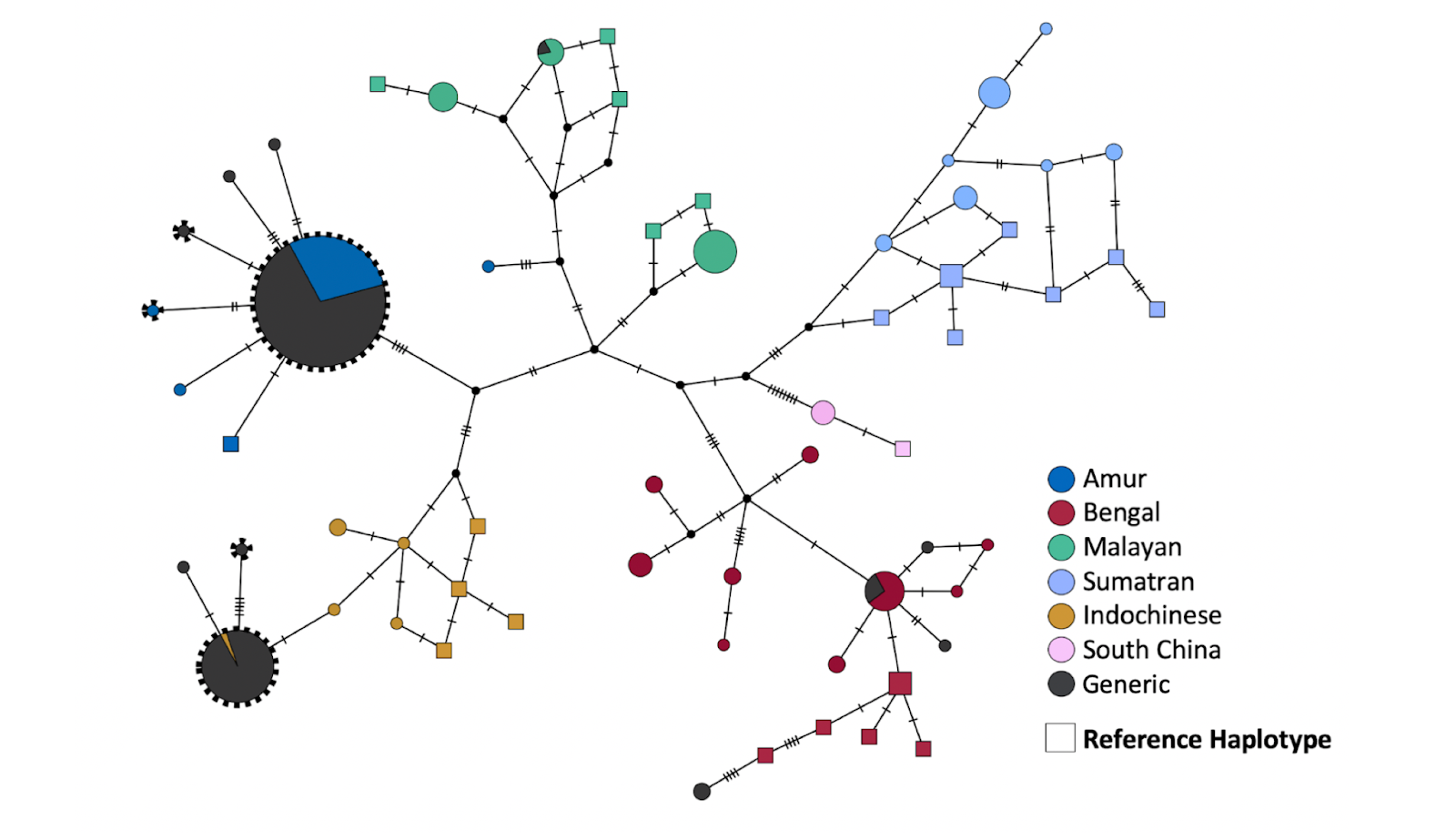

**Supplementary Fig.** **2** Median haplotype joining networks for all samples after numt contamination removal in any gene.

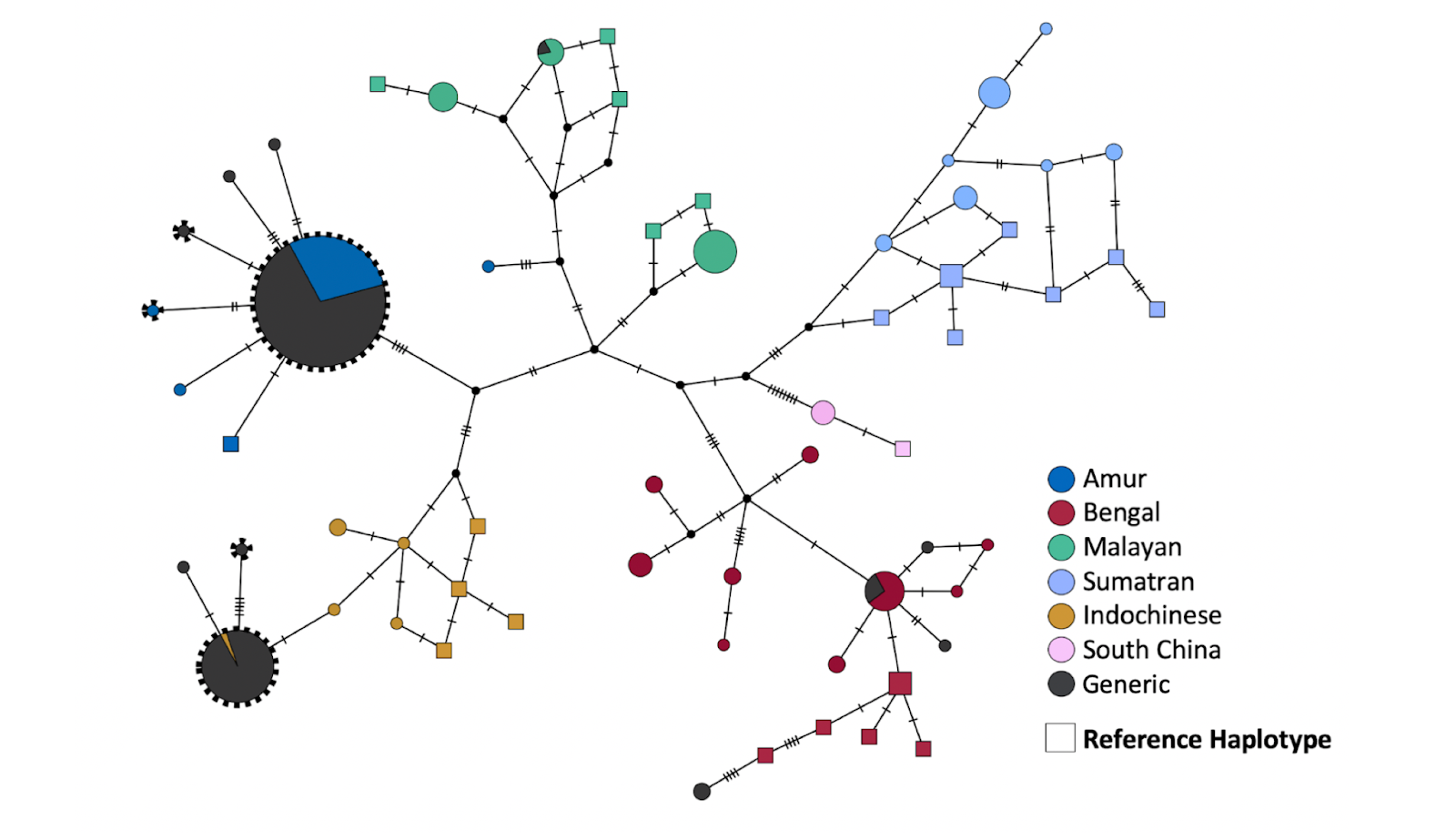

**Supplementary Fig.** **3** Median joining haplotype network based on 3,605bp of mitochondrial sequence for 273 samples, including 25 reference haplotypes from Luo et al. 2004. Each hatch mark represents a nucleotide change.

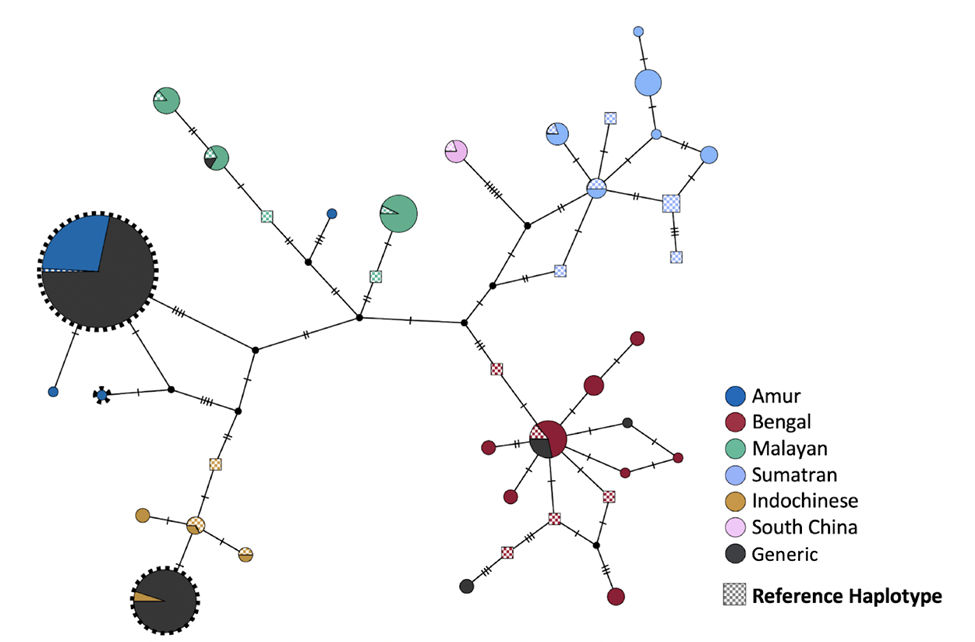

**Supplementary Fig.** **4** Median joining haplotype network based on 3,255bp of mitochondrial sequence for 274 samples, including 25 reference haplotypes from Luo et al. 2004. Gene 1 and Gene 2 sequence data are not included. Eleven samples that were removed in the haplotype map shown in Fig. 1 due to numt contamination of Gene 1 or Gene 2 are included in this network. The haplotypes represented in these eleven samples are indicated with a bold dotted outline.

#### Comparing local ancestry segments

In order to investigate local ancestry, we used the set of phased reference files generated for the imputation pipeline. Only individuals that were not duplicates were retained. Overall, the captive tigers contained primarily Amur ancestry, followed by Bengal ancestry, Sumatran ancestry, Malayan ancestry, and the least of the genomes came from Indochinese ancestry (Supplementary Fig. 5). Amur ancestry tracts also had the longest mean length (Amur 9,835,847bp; Bengal 5,843,489bp; Indochinese 2,095,711bp; Malayan 2,997,425bp; Sumatran 4,715,953bp). Only 34 local ancestry segments across any individual were longer than the median autosomal chromosome size (124,427,884), indicating that few tracts (if any) likely comprised entire chromosomes. Of course, since the reference genome used here contains gaps and since the real recombination map for tigers is unknown, these ancestry tracts are only approximate. Our ability to accurately detect local ancestry will improve with better reference genomes and larger reference databases.

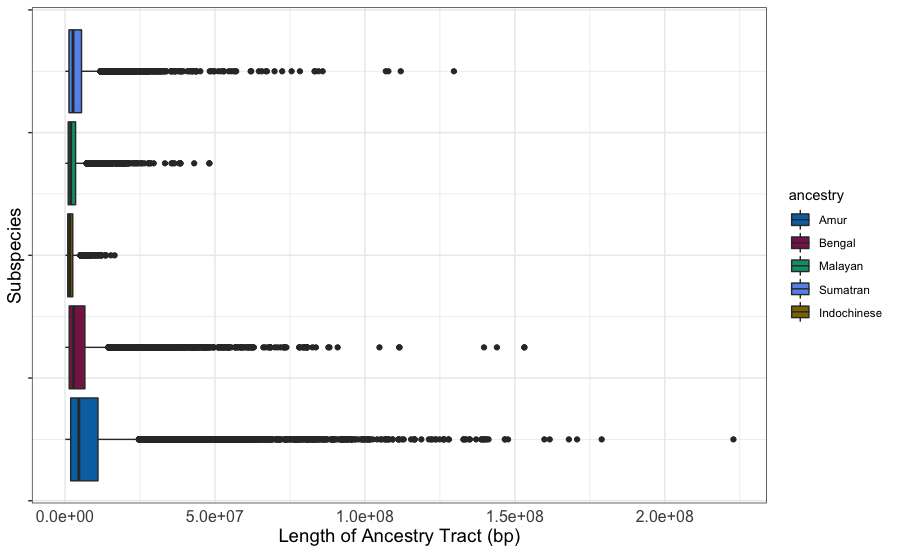

**Supplementary Fig. 5** Boxplot of ancestry tract sizes per species across all unimputed, phased individuals.

We next re-ran PCA (see *Supplementary Methods* for parameters) using only captive tigers. To determine whether the generic tigers showed signs of structure, we used PCA and Identity-by-state (IBS) clustering. We restricted to unimputed generic individuals only and ran PCA analyses in PLINK as previously described. We performed hierarchical clustering on the shared identity-by-state (IBS) loci between individuals to assess structure as well. The IBS matrix was made using the SNPRelate^30^ function ‘snpgdsIBS’ in R and hierarchical clustering was conducted with SNPRelate as well by calling the function ‘snpgdsHCluster’. Results from PCA and hierarchical clustering largely agree with each other. Using PCA, we did not observe any obvious structure (Supplementary Fig. 6A & 6B) and that nebulous clusters form in line with the top ancestry component of any individual. Hierarchical clustering supported the generics being classified as a single group as well (Supplementary Fig. 6C). Therefore, we can conclude that the structure of the generic population mimics the historical admixture from its founding and there are no distinct clusters formed by the various breeding facilities the tigers were taken from. Most likely, individuals are traded between facilities/locations often enough that the tigers form one, well-mixed population. For main figures, the outlier individual (EFRCT18) was removed. The outlier individual in the PCA that was removed (Supplementary Fig. 6 A, B) was found to have a unique ancestry signature in the population, having greater than 10% ancestry of all subspecies except South China, which was unique among individuals.

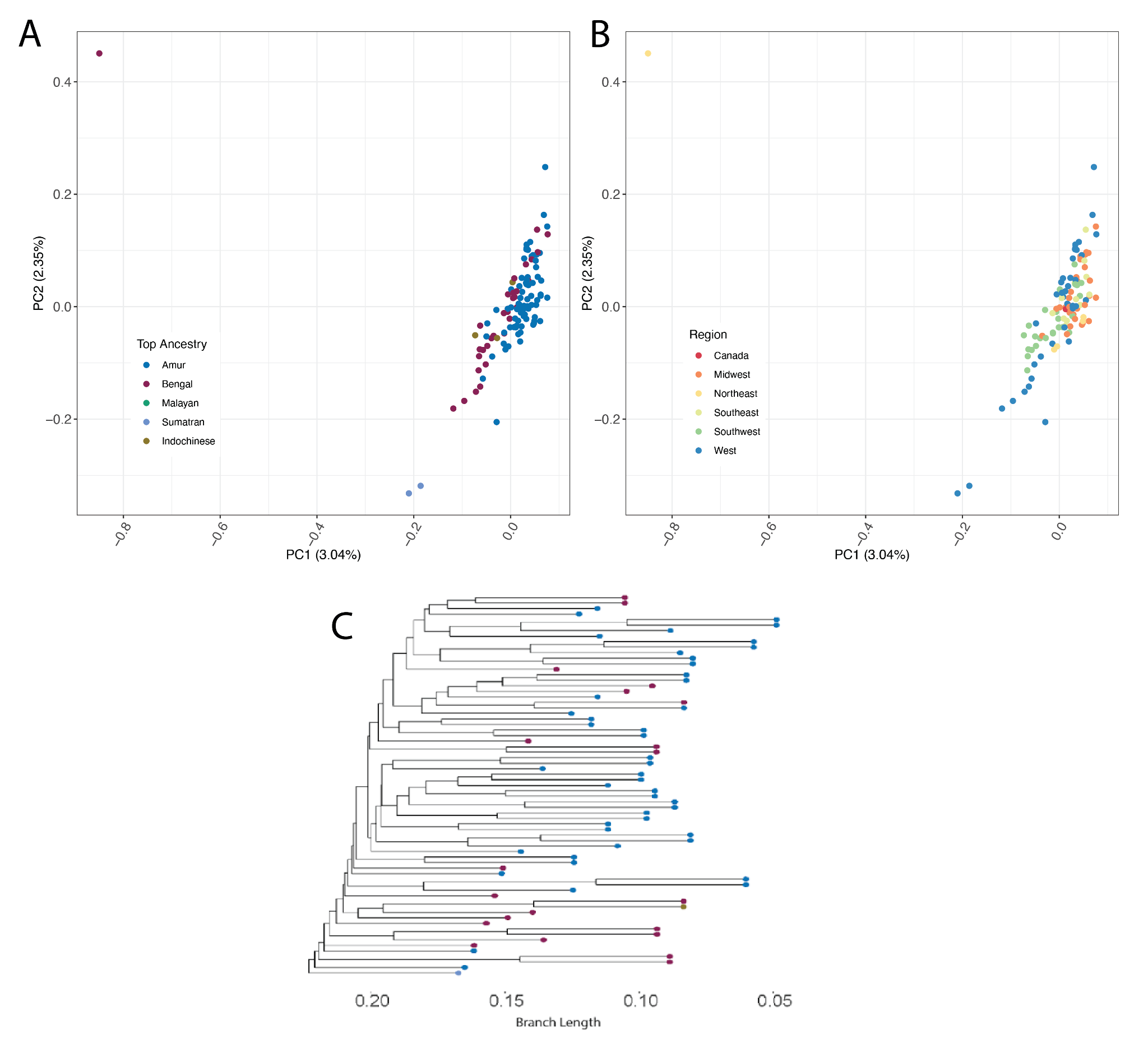

**Supplementary Fig.** **6** A) Principal component analysis (PCA) of autosomal sites for generic tiger colored by their top ancestry component. B) PCA of autosomal sites colored by their birthplace of origin. C) clustering based on identity-by-state (IBS) sharing between generic individuals. Individuals are labeled with their corrected subspecies designation.

#### Quantifying allelic diversity

We used the program ADZE v1.0^31^ to investigate how diversity was distributed across the various tiger groups. Using only unimputed, high-coverage individuals (>5x), we calculated both private allelic diversity and allelic richness. We did not include the South China tiger in these calculations, since we only had a single individual in the unimputed dataset. Due to the limited sample size of the Indochinese subspecies (N = 6), and since ADZE requires a holdout of two for the private variation analyses, we also ran the same analyses without the Indochinese tigers.

Analyses of allelic richness revealed that the Bengal tiger subspecies had the highest amount of allelic diversity (Supplementary Fig. 7 & 8), followed by the Generic and Malayan tigers. Sumatran and Amur tigers had less diversity overall. Indochinese tigers appear to have a comparable diversity to the Bengal tigers, but because of the reduced sample size, it is unclear whether additional individuals would place them above or below the Bengal tiger group. Analyses of the private allelic diversity showed that despite having high amounts of diversity, the generic tigers contain very few private alleles compared to most other subspecies, reflecting the admixture in their genomes (Supplementary Fig. 7 & 8). The Amur tiger subspecies had the fewest private alleles, and the low allelic richness and lack of private variation in this group suggests a history of severe bottlenecking. Bengal tigers had by far the most amount of private variation. Despite these results, it is clear from the plots that more samples from each group would benefit our understanding of the shared variation and history of these groups.

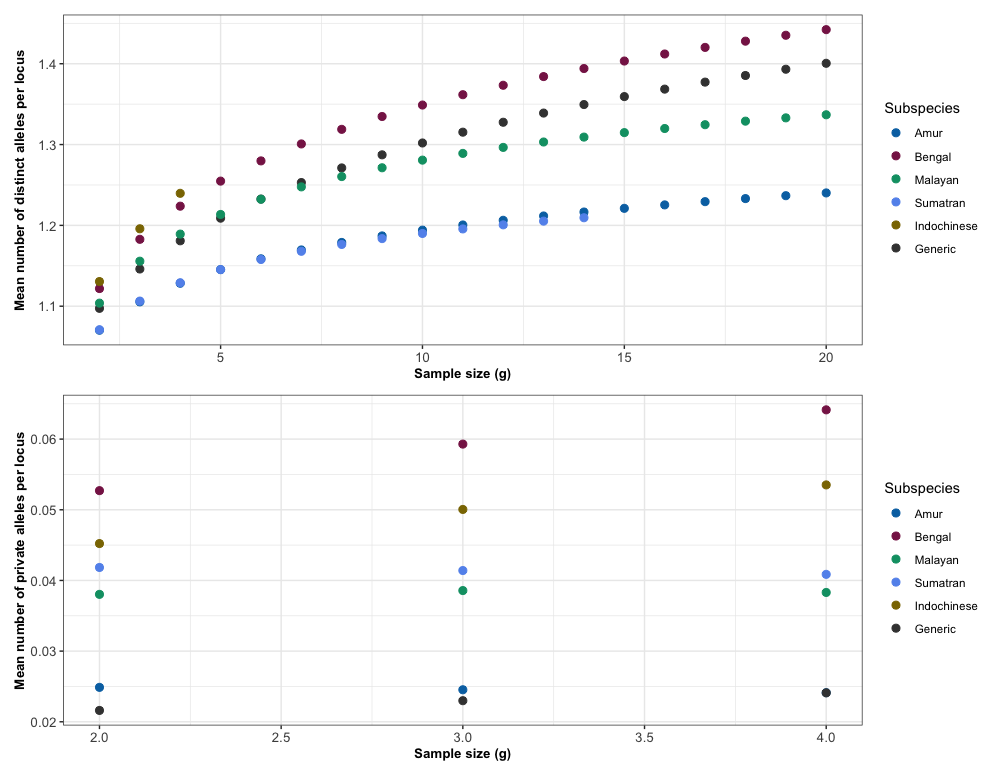

**Supplementary Fig. 7** (Top panel) ADZE analyses showing the mean allelic richness per group as sample size increases. (Bottom panel) ADZE analyses showing the mean number of private alleles per locus per group as sample size increases. This figure includes the Indochinese.

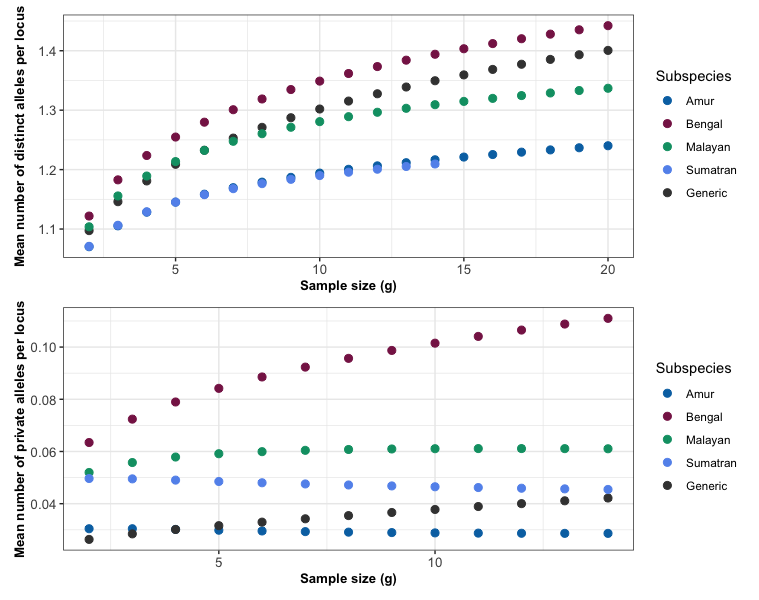

**Supplementary Fig. 8** (Top panel) ADZE analyses showing the mean allelic richness per group as sample size increases. (Bottom panel) ADZE analyses showing the mean number of private alleles per locus per group as sample size increases. Comparable to Supplementary Fig. 7, but here we have dropped the Indochinese to increase sample size.

#### Pairwise sharing of IBD segments

IBD segments were identified separately for each subspecies using TRUFFLE v1.38 ^20^, on only unimputed and unrelated individuals that had greater than 5× coverage. Since we only used the unimputed individuals, we were once again forced to drop the South China subspecies from this analysis. The pairwise sharing between each pair of unrelated individuals is displayed in Supplementary Fig. 9.

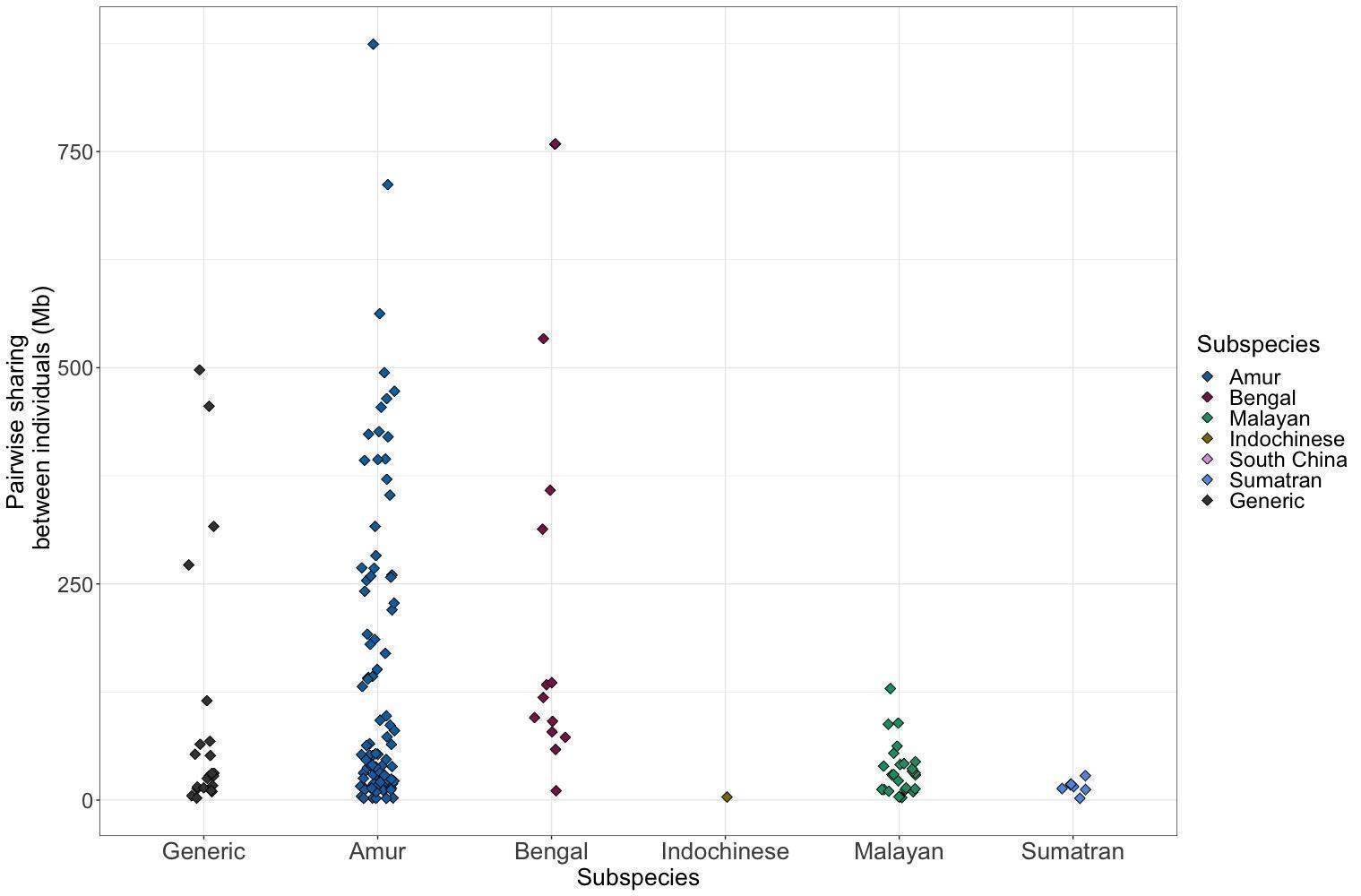

**Supplementary Fig.** **9** IBD sharing between pairs of unimputed samples in each subspecies. We observed increased sharing in the Amur and Bengal subspecies relative to Indochinese, Malayan, and Sumatran subspecies.

IBD scores and fold enrichment of wild subspecies relative to the generic population can be seen in Supplementary Fig. 10.

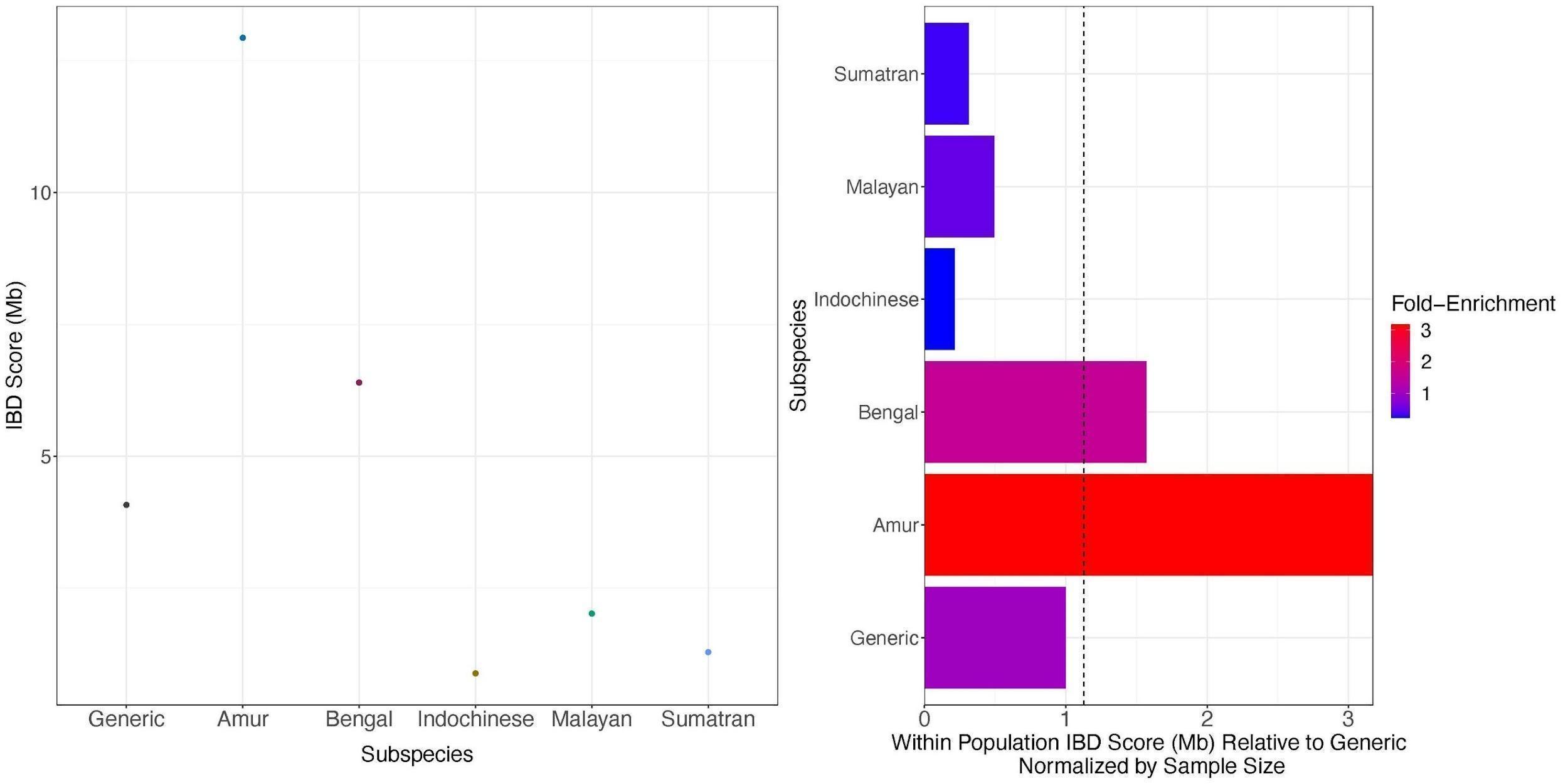

**Supplementary Fig.** **10** IBD for each subspecies of wild tiger and captive tigers. A) The IBD score is the amount of pairwise sharing between unrelated individuals. We can see that the largest IBD score is seen in the Amur subspecies followed by the Bengals then captives. B) Fold-enrichment of IBD in each subspecies relative to captive population. We see a large enrichment of IBD in the Amur and Bengal subspecies relative to the captives. Conversely, we see depletions of IBD sharing in the Sumatran, Malayan, and Indochinese subspecies relative to the captive population.

#### Site frequency spectrum

Site frequency spectra (SFS) were generated using only unrelated and unimputed samples with greater than 5× coverage. Since we only used the unimputed individuals, we were forced to drop the South China subspecies from this analysis. We chose two groups of samples, which included ten and six unrelated individuals from each subspecies. We chose these numbers so that we could include the Indochinese subspecies in our analyses. The six unrelated individuals are a subset of the ten unrelated individuals. Supplementary Fig. 11 contains all four SFS for wild and captive tigers. Generic tigers have the largest fraction of singleton variants followed by the Bengal tigers. The Sumatran tigers have the largest fraction of high frequency derived sites followed by the Amur tigers.

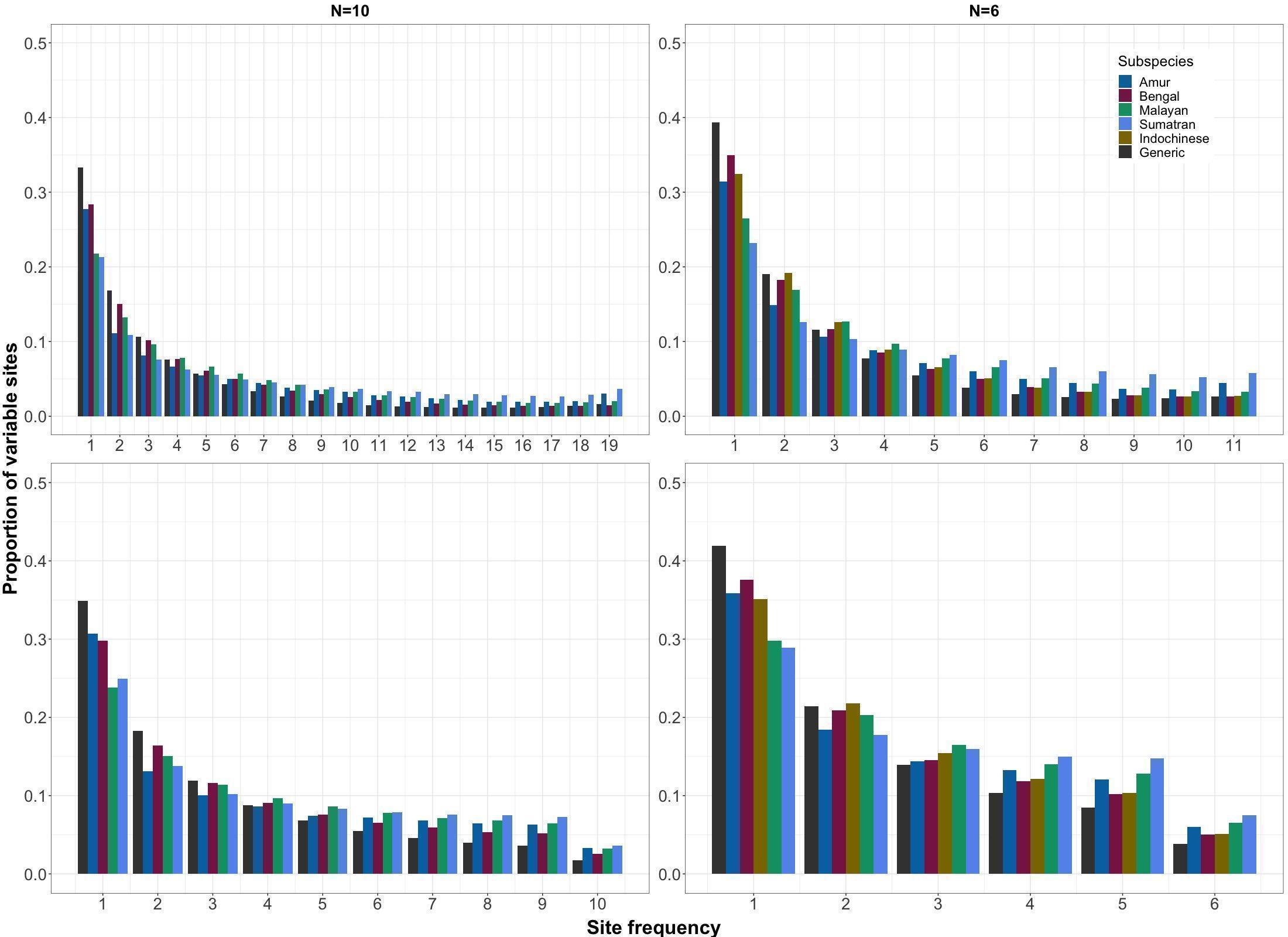

**Supplementary Fig.** **11** Folded and unfolded site frequency spectra for N=10 and N=6 individuals. The top panel is the unfolded SFS, and the bottom panel is the folded SFS. Variants were polarized using the Progressive CACTUS ancestral base.

#### Computing Genetic Load

We annotated sites in our VCF with VEP and SIFT annotations (*see Supplementary Methods*). We used only unimputed individuals with coverage greater than 5×. Since we also had the ancestral allele from Progressive Cactus (*see SFS section*), we used multiple approaches to count deleterious variants in the genome of each unimputed individual: 1) tabulating homozygous derived genotypes (counting homozygotes); 2) counting the total number of homozygous and heterozygous derived genotypes (counting variants); and 3) summing twice the number of homozygous derived genotypes plus heterozygous genotypes (counting alleles). If deleterious alleles act recessively, then counting derived homozygotes is most relevant to disease.  If deleterious alleles are recessive, counting derived homozygotes is most relevant, and counting alleles is most relevant when deleterious alleles have additive effects on fitness ^32,33^. The deleterious and neutral variation contained in Fig. 4 in the main text and Supplementary Figs. 12 & 13 mirror each other except that Supplementary Fig. 12 contains all counting methods and Supplementary Fig. 13 contains outlier individuals (GEN1 and BEN_NE2). Additionally, we re-did counts with synonymous (SYN) and nonsynonymous (NS) variation. We saw the same pattern except that there are more variants that are annotated as either SYN or NS than putatively neutral or putatively deleterious. Lastly, we found that individuals with the most putatively deleterious derived homozygotes also tend to have the largest inbreeding coefficients, quantified with either F_SNP_ or F_ROH_ (Supplementary Fig. 14).

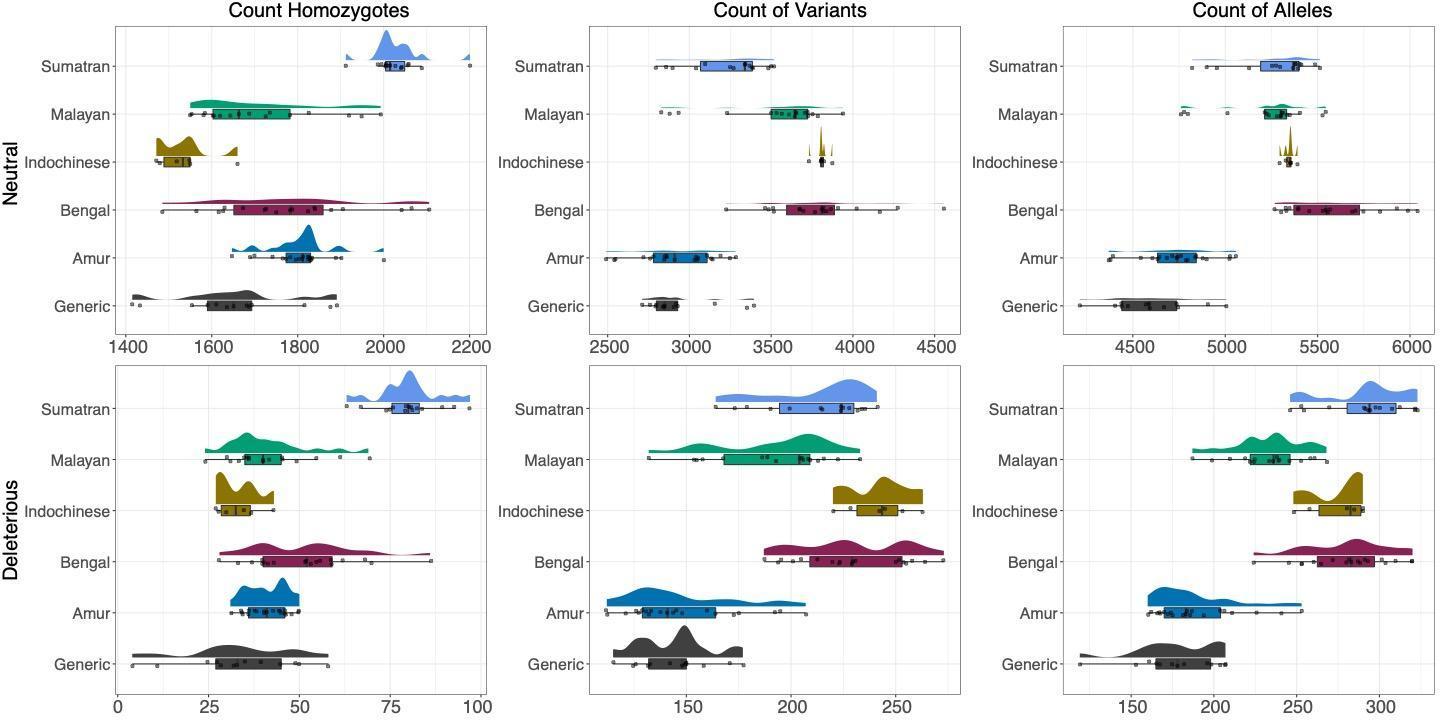

**Supplementary Fig.** **12** (Top row): Neutral variation in each tiger subspecies and generic tigers using different models. (Bottom row): Deleterious variation in each tiger subspecies and generic tigers using different models. Count homozygotes represents only homozygous deleterious variation; count variants represents both deleterious homozygotes and heterozygotes equally (both count for one deleterious variant); and count alleles weights homozygotes as two and heterozygotes as one.

Interestingly, there were two individuals, one each in the generic and Bengal populations, which were outliers in terms of both the counting variants and alleles analyses (Supplementary Fig. 13). Neither of these individuals were outliers in any other analysis we conducted, despite having an almost 3-fold enrichment of heterozygous sites that were annotated. The enrichment of heterozygous sites was validated via examination of the read counts for the reference and alternative alleles in these individuals. We believe our results capture one of the pitfalls of applying annotations from one species (cat) to another (tiger), in which subsets of sites in the genome are annotated and not necessarily representative of the full spectra of possible mutations, due to mismatches that occur during liftOver, causing some sites to be lost. Our results should caution other researchers who are attempting similar analyses.

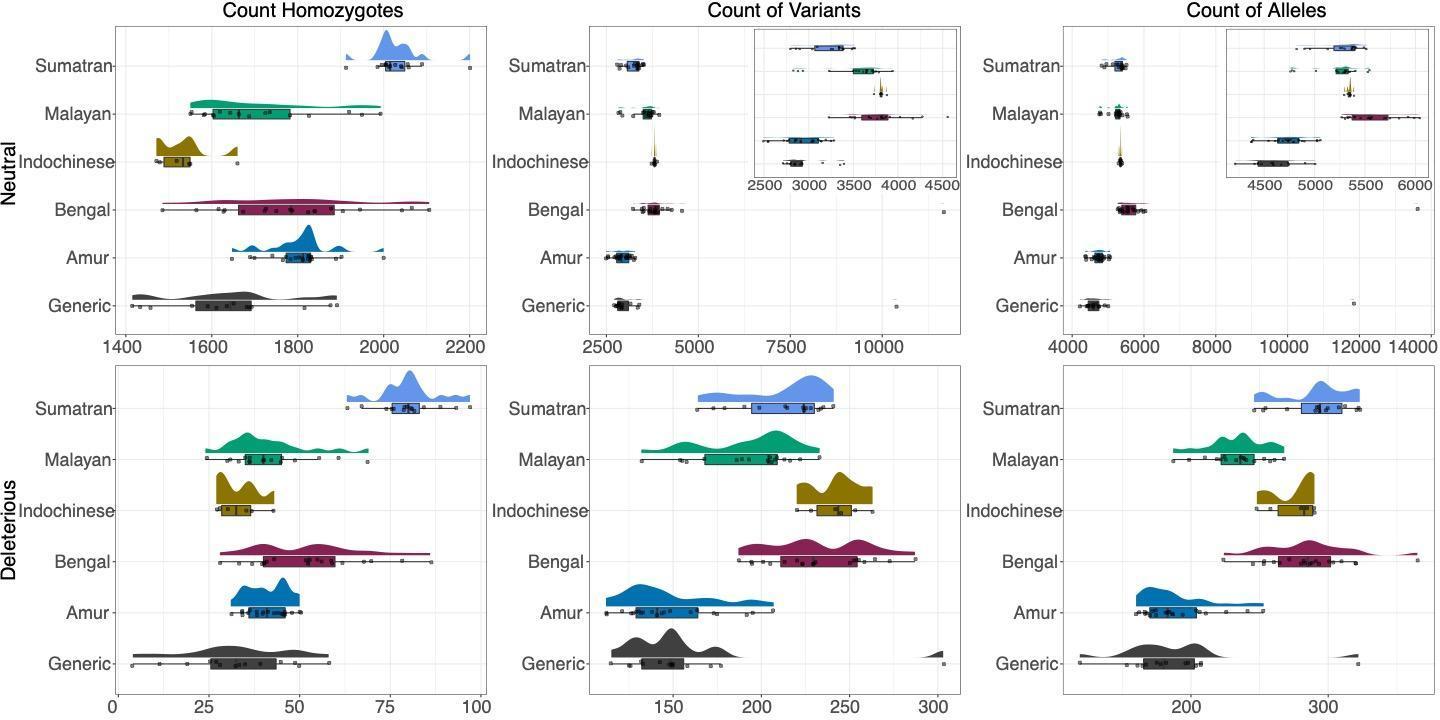

**Supplementary Fig. 13** Neutral versus deleterious counts of homozygotes, variants, and alleles. The inset zooms in on the non-outlier portion of the graph. Outlier individuals are GEN1 and BEN_NE2. Count homozygotes represents only homozygous deleterious variation; count variants represents both deleterious homozygotes and heterozygotes equally (both count for one deleterious variant); and count alleles weights homozygotes as two and heterozygotes as one.

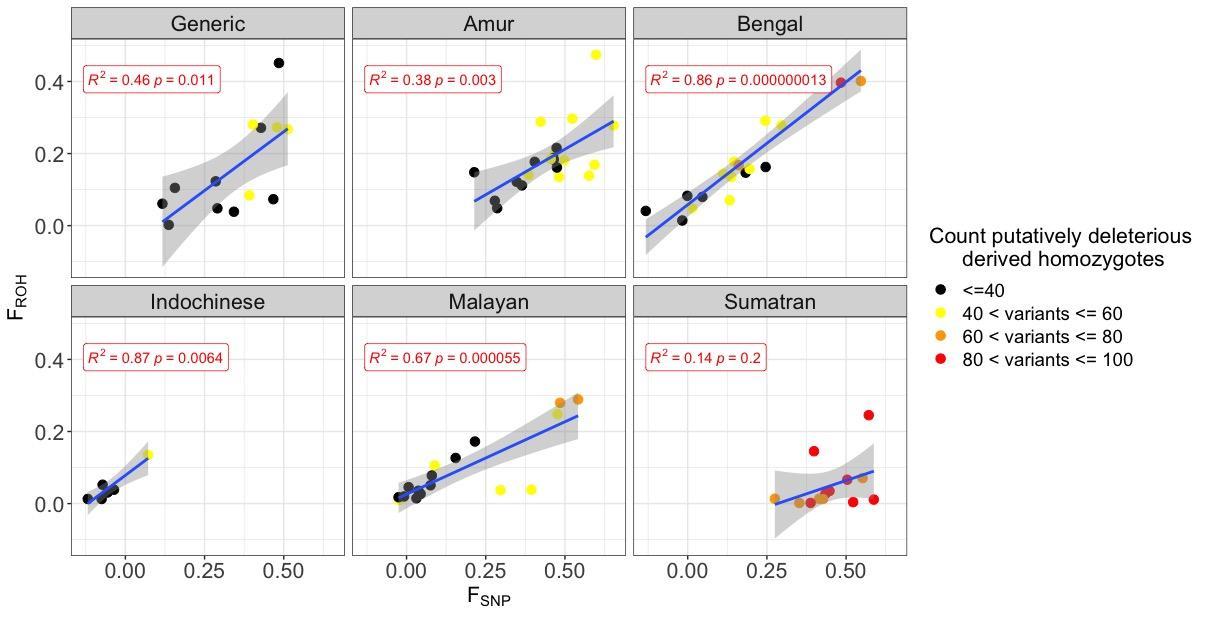
**Supplementary Fig. S14:** Pearson correlation between F_SNP_ (x-axis) and F_ROH_ (y-axis) for each subspecies. Individuals are labelled with the number of putatively deleterious derived homozygotes in their genome.

#### Quantifying the enrichment of nonsynonymous and deleterious variation within ROH

We tested whether there is an enrichment of nonsynonymous or putatively deleterious mutations in ROH over non-ROH regions for the three different ways of counting variation. To account for differences in neutral variation, we standardized by synonymous or putatively neutral variation. Then, we calculated the ratio of nonsynonymous over synonymous variation in ROH regions divided by the ratio of nonsynonymous over synonymous variation outside of ROH. We computed significance by generating a contingency table and running fisher.test() in R. We repeated the analysis for putatively deleterious and putatively neutral variation within and outside of ROH.

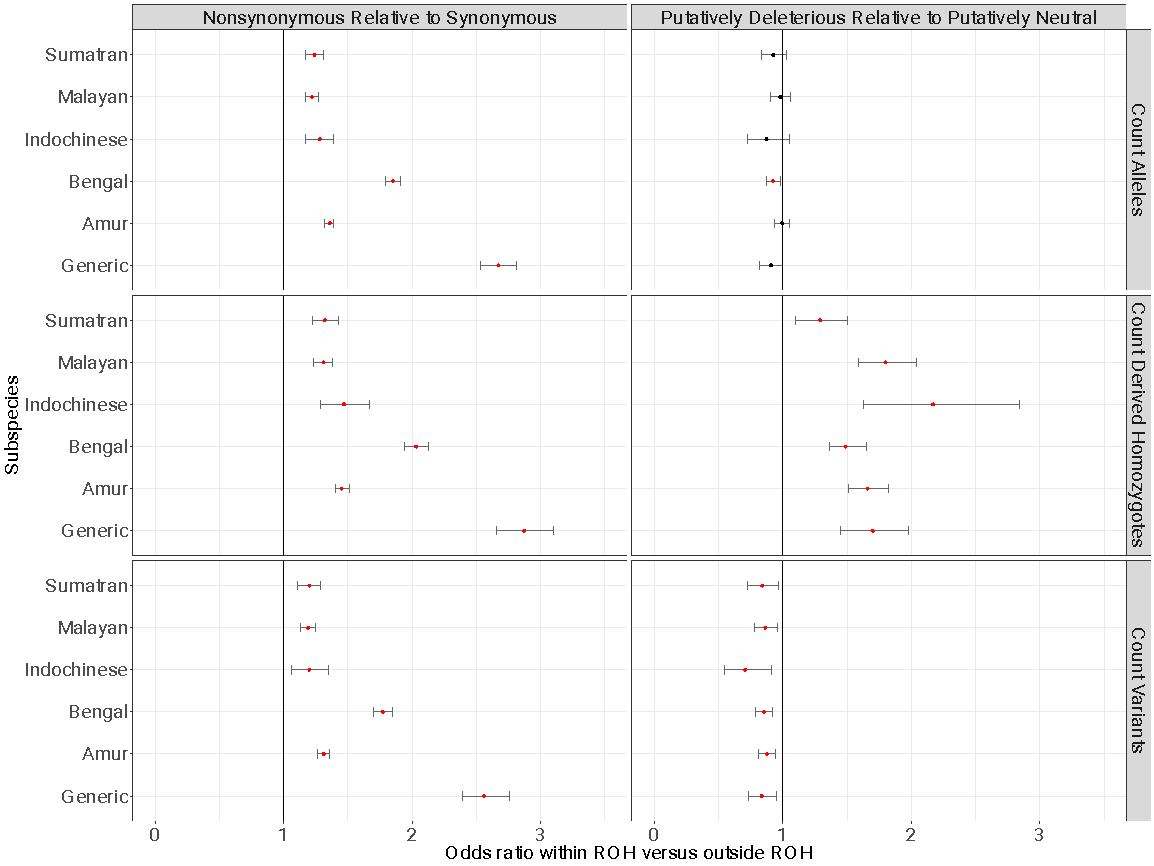

**Supplementary Fig. S15:** Odds ratio of variation falling within or outside of an ROH (x-axis) and Subspecies of interest and counting method (y-axis). If the p-value (Table S2) is significant the dot is filled in red. The left column is nonsynonymous variation relative to synonymous and the right column is putatively deleterious relative to putatively neutral. Generic tigers are a clear standout in the case of nonsynonymous variation. However, all populations are similar in the case of putatively deleterious variation.

#### Concordance and accuracy of imputation pipeline

To examine the accuracy and utility of our imputation pipeline, we investigated the concordance of variant calls across different depths. Additionally, a primary purpose of building the imputation pipeline was to accurately call ancestry and identify individuals in low-coverage and unknown samples, so we also examined the accuracy of these measures across samples with different depths.

We examined individuals from both the imputed and unimputed sample sets. For each of these individuals, we down-sampled reads to approximately 5×, 2×, 1×, 0.5×, and 0.25×. We then input these down sampled files into the Gencove pipeline for imputation. The resulting VCFs were restricted to the high-quality sites identified in Supplementary section 1.2.2 using BCFtools *view* -R. We then compared the calls from each of these to the calls in the original reference file using vcf-compare from VCFtools with the flag ‘-g’, which in addition to comparing which sites are present, compares the actual genotype calls. As expected, we found that with increasing depth, the non-reference discordance rate (NDR) decreased (Supplementary Table S3) in all but one individual (MAL1). In general, NDR remained fairly low even at the lowest depth (0.25×) we tested for individuals in the reference data set (7.71-14.79%; Table S3). For imputed individuals that were not included in the reference panel, we found that, compared to the raw GATK calls, imputation performed comparatively with higher coverage individuals, but the NDR increased much more drastically with decreasing coverage. This was especially true for the individual with South China ancestry that we tested (SRR7651468) since we only had one individual representative of this population in the reference dataset. Naturally, this demonstrates that reference panels are drastically improved with more representative individuals.

To quantitatively compare the ancestry calls with down-sampled and imputed data, we created a distribution of each ancestry category (Amur, Bengal, Indochinese, Malayan, South China, and Sumatran) composed of the assigned ancestry proportion from each of the nine tested individuals. We then tested whether there was a significant difference in the distributions of assigned ancestry calls between the down-sampled data relative to the ancestry inferred without imputation or down-sampling using a Kolmogorov–Smirnov test with the ks.test() function in R.

Despite the variation in NDR across imputed and unimputed samples using the imputation pipeline, the predicted ancestry of down sampled individuals remained fairly accurate across coverages (Supplementary Fig. S16). For individuals that were verified as single-subspecies ancestry, the ancestry inference remained at or close to 100% across all coverages. This indicates that the imputation panel can accurately assign relative ancestry components even at ultra-low (0.25×) coverages. We tested relatedness estimate accuracy for a subset of samples (two imputed, two unimputed) by examining the similarity of individuals detected as related above a certain threshold to the sample in question. To summarize these results, we tabulated only relatedness values over 0.177 (~second degree relatives) for the individual in question.

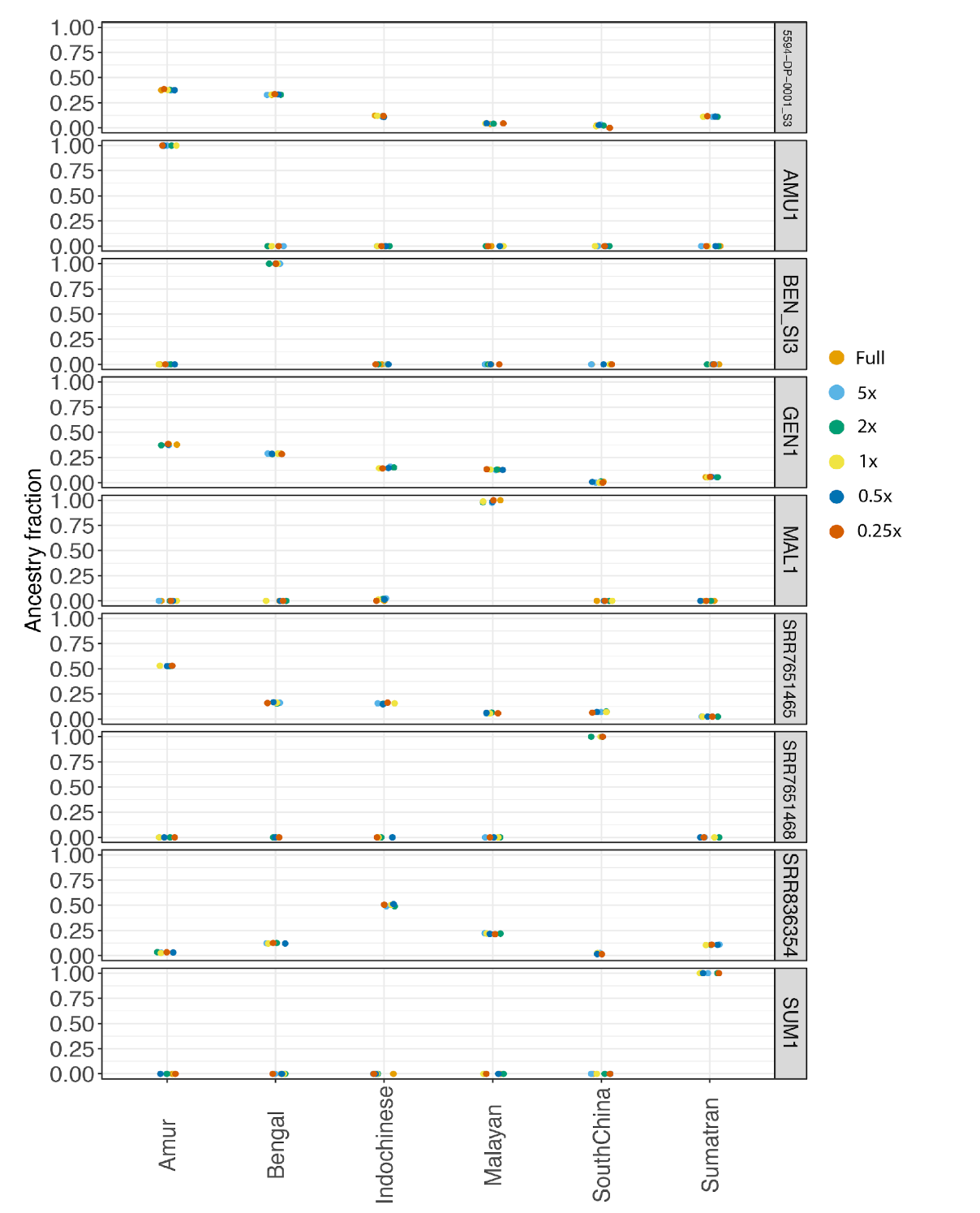

**Supplementary Fig. S16:** Relative ancestry components during imputation across various coverage thresholds across nine individuals with different ancestries.

We found that of the four individuals tested, all samples were able to be identified as the same sample (meaning individuals were able to be identified irrespective of depth, Supplementary Table 5). In addition, all first-degree relatives detected in the original dataset were also identified as first-degree relatives when the data was subsampled for unimputed individuals (Supplementary Table 5). However, the accuracy of kinship estimates declined (compared to those using the full set of data) for imputed individuals as depth decreased. Though it is outside of the scope of this study, the accuracy of these estimates should be assessed more carefully using sample sets with additional known relatives.

#### Ancestry verification and duplicate removal

To verify the ancestry of the tigers, we first used PCA. PCA first confirmed that the designated subspecies in the unimputed dataset all formed unique clusters (Supplementary Fig. 16). Although we only had a single individual from South China, we still observed this individual to be separate from all the other clusters in PCA space across all principal components that we examined (Supplementary Fig. 16). Further, this individual has previously been confirmed as having a distinct mitochondrial genome^34^. The South China tiger lineage is functionally extinct, and the remaining captive population was founded from just six individuals in the 1950s and 1960s. Previous studies have suggested that the lineage was mixed with at least the Indochinese and possibly the Amur subspecies^35,36^. Given this information and the fact that new studies with additional South China individuals have confirmed their uniqueness and mitochondrial placement^37^, we opted to use this individual in the reference set, despite its potential admixture. With so few individuals, we felt that this reference individual was representative of the extant South China population and further sequences can be added to the reference database when they are available.

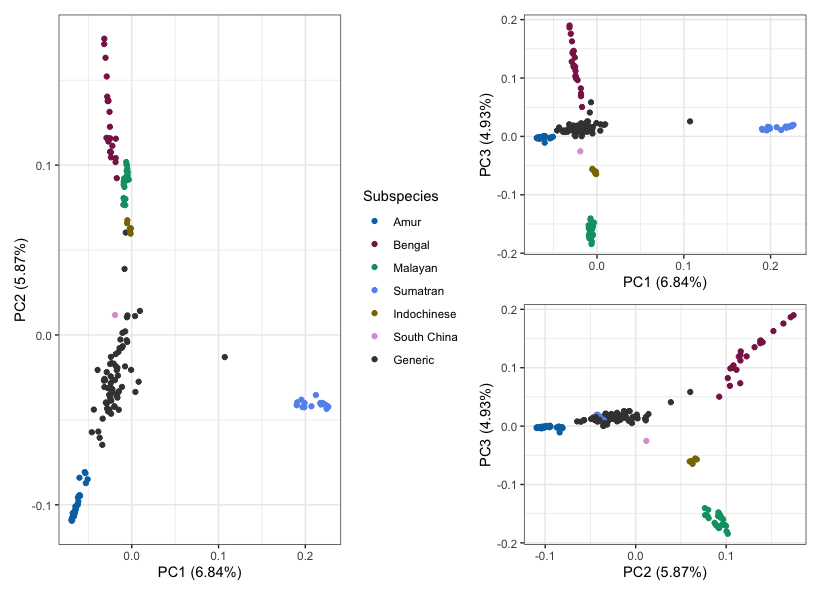

**Supplementary Fig. 16**: PCA of all unimputed individuals.

PCA revealed a number of individuals that were likely misidentified from the imputed sample set (Supplementary Fig. 17). As a result, we relabeled the population assignment of six individuals to ‘generic’ after verifying that they were admixed (see below). Five individuals (SRR7651464, SRR7651465, SRR7651466, SRR7651467, SRR7651470) were originally labeled as Amur and one individual (SRR836354) that was originally labeled as a Bengal tiger (Supplementary Fig. S17 & S18).

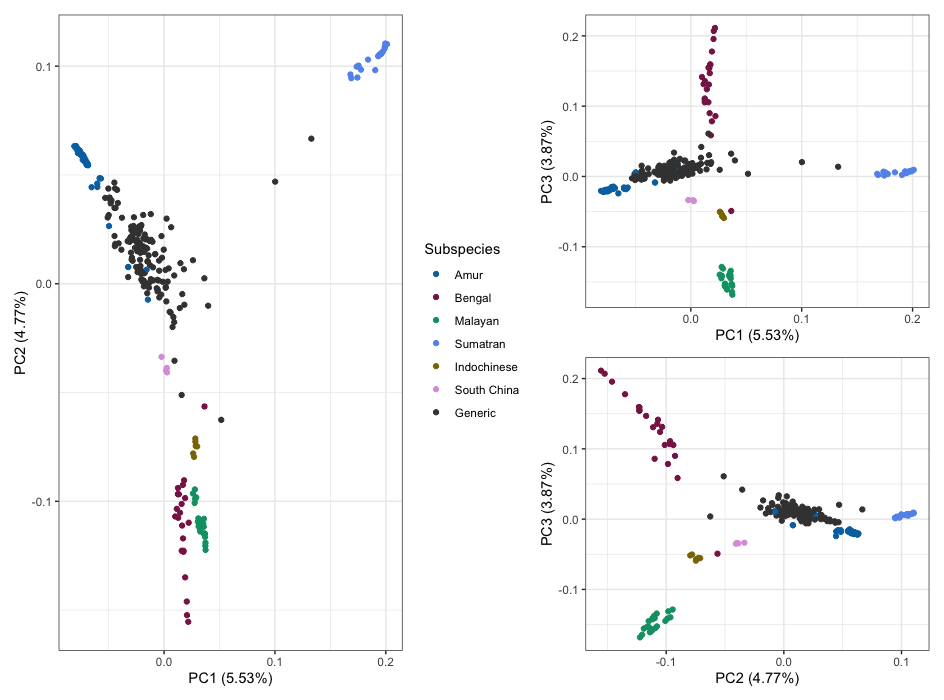

**Supplementary Fig. 17**: PCA of all individuals (unimputed and imputed) prior to duplicate removal or ancestry correction.

To investigate ancestry fractions across all individuals, we used the program ADMIXTURE v1.3.023^38^. All individuals of verified single subspecies ancestry (see above) in the unimputed dataset were used as reference individuals according to their assigned subspecies. Tigers in the imputed dataset and individuals of unknown ancestry in the unimputed dataset were then evaluated using a supervised analysis, with otherwise standard parameters (Supplementary Fig. 17).

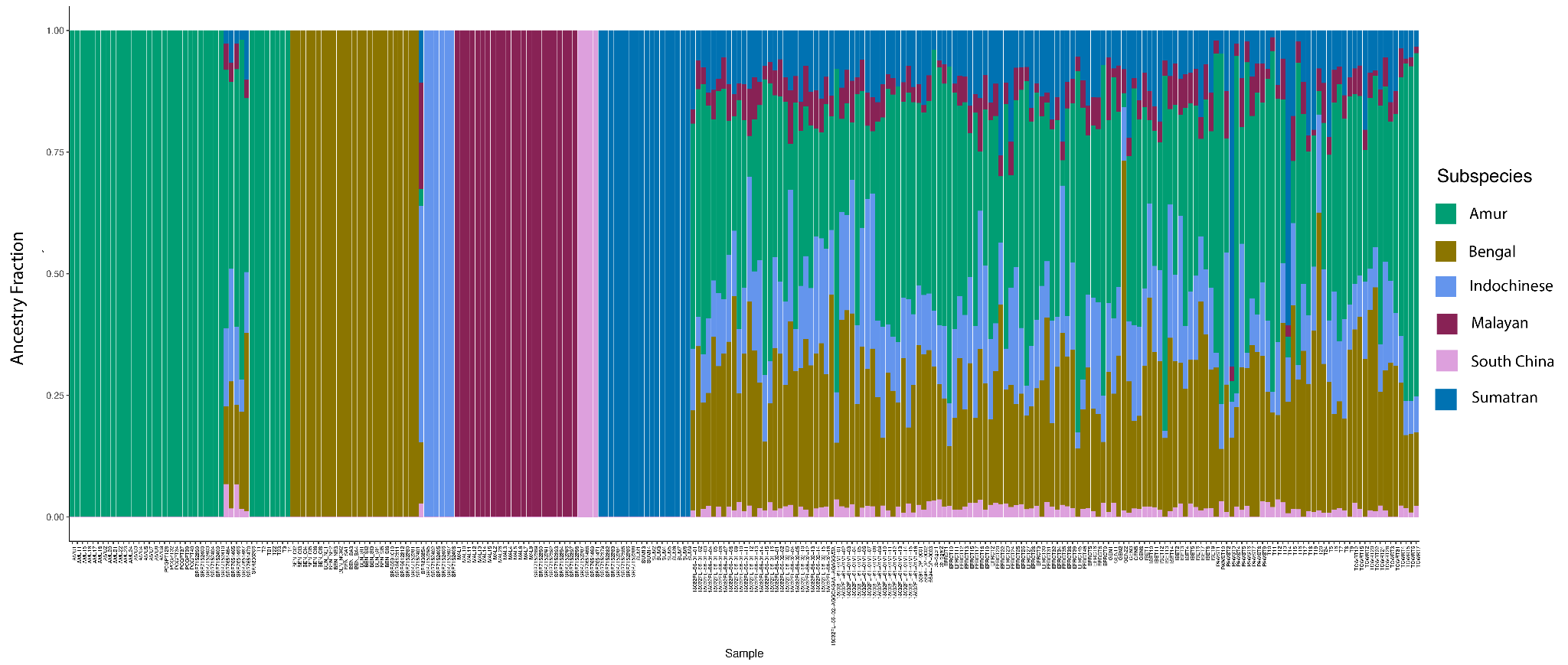
**Supplementary Fig. 18** Admixture plot of all individuals (unimputed and imputed) prior to duplicate removal. Individuals grouped according to the original ancestry record.

Based on results in Section 1.2.4, we next ran VCFtools^7^ and SNPRelate^30^ to profile relatedness. IBDMLE within SNPRelate did slightly better overall than VCFtools, but this was not consistent across populations (Supplementary Fig. 19A). VCFtools and IBDMLE identified three and two pairs of individuals that were potential duplicates, respectively, one of which overlapped (SRR7651464, SRR7651466). Upon further investigation, we found that the second individual identified by IBDMLE was indeed a duplicate due to two different spellings of the sample (EFRCT6, Sampson; EFRCT8, Samson), but that the additional two individuals identified by VCFtools had no other evidence of being duplicates. As a result, we only identified duplicated individuals using IBDMLE scores (SRR7651466 and ERCT8 were removed).

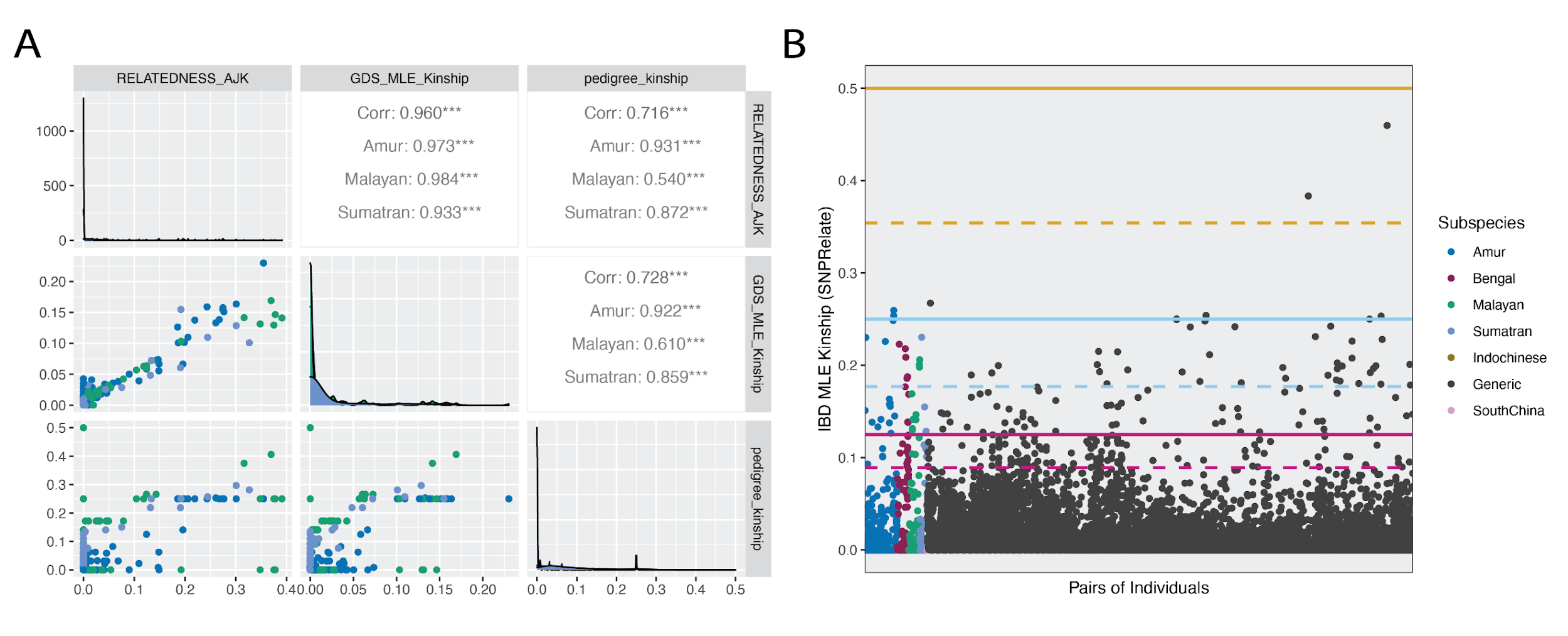

**Supplementary Fig. 19** A) Correlations of pedigree relatedness to kinship as estimated by VCFtools relatedness and SNPRelate IBDMLE including low coverage samples. B) Distribution of estimated kinship using IBDMLE. Lines represent first degree (yellow), second degree (light blue) and third degree (maroon) relative lines. Dotted lines represent the geometric mean for each estimate.

#### Heterozygosity and missing data

We investigated the heterozygosity of the various populations using VCFtools ‘--het’. Heterozygosity was calculated by dividing the observed heterozygosity (OHOM) from the output with the number of callable sites. The number of callable sites was calculated by subtracting the number of sites filtered for mappability from the total number of autosomal sites.

Since we observed clustering of imputed heterozygosity values (Supplementary Fig. 20), we concluded that these values were not reliable. We additionally tested to see if heterozygosity was correlated to the percentage of missing sites. Using VCFtools, we calculate the proportion of missing sites per individual using VCFtools ‘--missing-indv’.

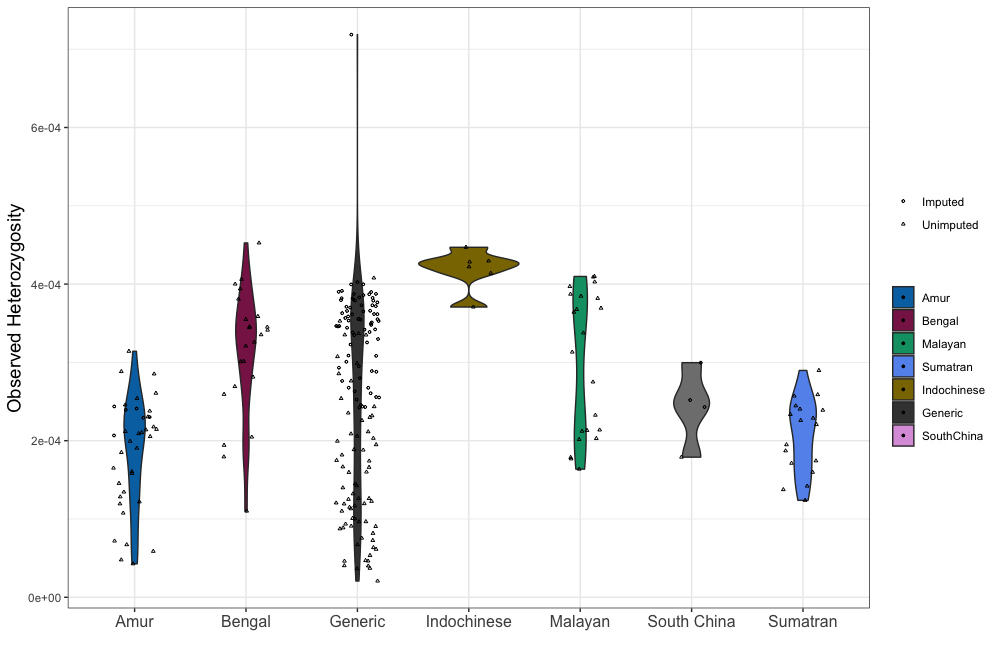

**Supplementary Fig. 20** Observed heterozygosity as calculated by VCFtools for all samples without duplicates.

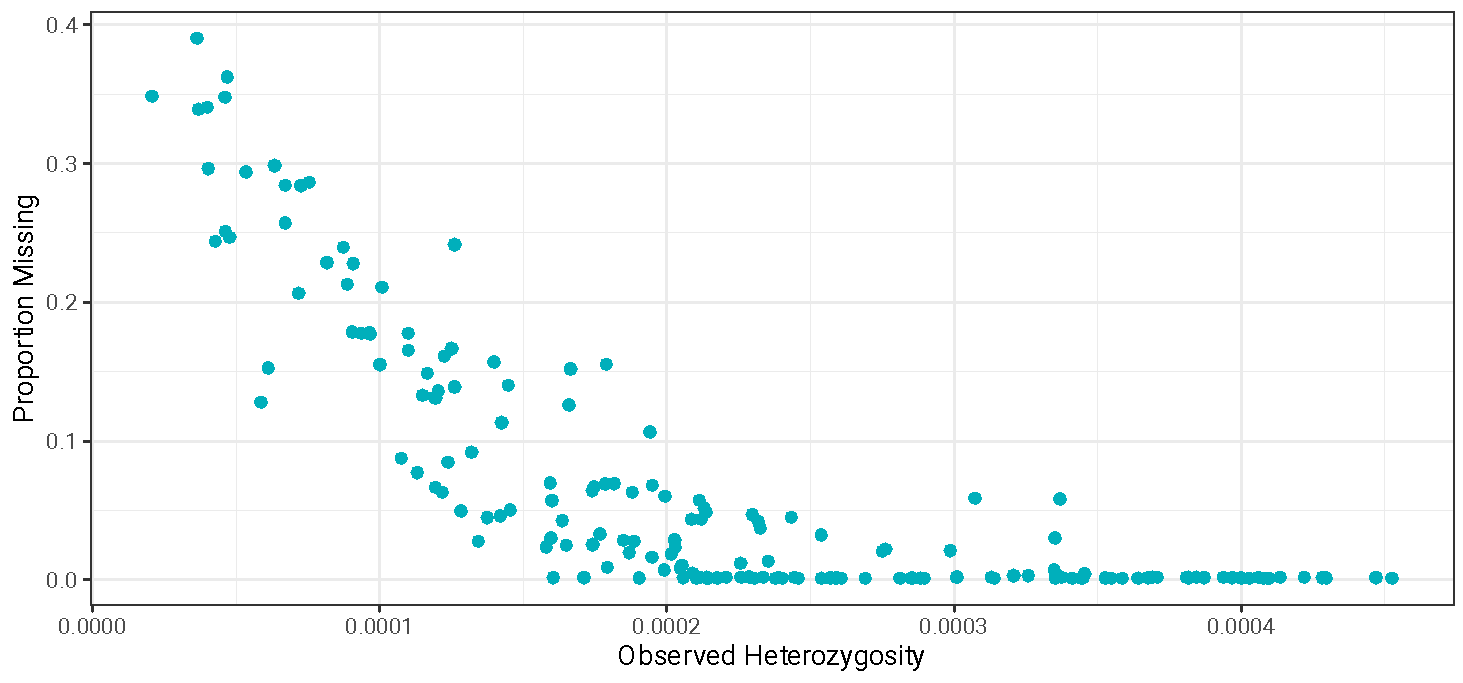

**Supplementary Fig. 21** Scatter plot showing the correlation of observed heterozygosity to the proportion of missing data per individual.

We found that heterozygosity appeared to be correlated with missingness (Supplementary Fig. 21), where data with more missing sites had lower observed heterozygosity. We thus decided to remove individuals with more than 20% of data missing to calculate the observed heterozygosity for each group, to conserve as many data points as possible, but also exclude as many outliers as possible.
